# E4F1 and ZNF148 are transcriptional activators of the A57C and wildtype *TERT* promoter

**DOI:** 10.1101/2023.01.13.523884

**Authors:** Boon Haow Chua, Laure Ferry, Cecilia Domrane, Nurkaiyisah Zaal Anuar, Anna Wittek, Sudhakar Jha, Falk Butter, Daniel G. Tenen, Pierre-Antoine Defossez, Dennis Kappei

## Abstract

Point mutations within the *TERT* promoter are the most recurrent somatic non-coding mutations identified across different cancer types, including glioblastoma, melanoma, hepatocellular carcinoma, and bladder cancer. They are most abundant at C146T and C124T and rarer at A57C, with the latter originally described as a familial case but subsequently shown also to occur somatically. All three mutations create *de novo* ETS (E-twenty-six specific) binding sites and result in the reactivation of the *TERT* gene, allowing cancer cells to achieve replicative immortality. Here, we employed a systematic proteomics screen to identify transcription factors preferentially binding to the C146T, C124T and A57C mutations. While we confirmed binding of multiple ETS factors to the mutant C146T and C124T sequences, we identified E4F1 as an A57C-specific binder and ZNF148 as a *TERT* WT binder that is excluded from the *TERT* promoter by the C124T allele. Both proteins are activating transcription factors that bind specifically to the A57C and wildtype (at position 124) *TERT* promoter sequence in corresponding cell lines and upregulate *TERT* transcription and telomerase activity. Our work describes new regulators of TERT gene expression with possible roles in cancer.

## INTRODUCTION

The breakthrough discovery of recurrent somatic point mutations in the *TERT* (telomerase reverse transcriptase) promoter – namely the C146T, C124T and A57C mutations (positions relative to the ATG start codon)(1, 2) – has highlighted the significance of non-coding mutations in the process of carcinogenesis. This exemplifies how creation or destruction of a transcription factor binding site in regulatory elements can contribute to carcinogenesis and tumor progression through upregulation of an oncogene or downregulation of a tumor suppressor gene. The *TERT* promoter mutations (TPMs) are generally mutually exclusive and occur in more than 50% of bladder cancer, adult glioblastoma, hepatocellular carcinoma (HCC) and melanoma (1–6), roughly at par with the various p53 mutations as the most frequently mutated protein-coding gene (7). Most of the point mutations take place at C146T and C124T, followed by A57C mutations. A57C was first identified in a family with a high incidence of cutaneous melanoma (1) and was later found to also occur somatically (8). Multiple other rarer variants have also been reported, which include tandem CC>TT conversions at positions −138/−139 bp and −124/−125 bp (0.8-10%) (1, 2, 8, 9), as well as point mutants G141T, C125A, C124A, C72G/C77T, C54A and G45T (all ∼1%) (5, 8, 9). Except for C125A and C124A, all mutations result in the creation of a *de novo* ETS (E-twenty-six specific) site. Reporter assays showed an increase in promoter activity for the C146T, C124T and A57C mutants as compared to the wildtype promoter sequence (1, 2, 10). The subsequent reactivation of the *TERT* gene, which encodes for the catalytic component of telomerase, is associated with increased telomerase activity (4). Unlike other core components of the telomerase enzyme, hTR (telomerase RNA) and dyskerin (DKC1), that are expressed ubiquitously (11), *TERT* expression is normally repressed in somatic cells. The majority (85-90%) of cancers maintain their telomeres by reactivating *TERT* expression, which contributes to their indefinite proliferative potential with critically shortened telomeres being a rate-limiting step in oncogenesis (12). Telomeres are composed of tandem TTAGGG repeats and exist as protective nucleoprotein structures at the ends of linear chromosomes (13, 14). During each cell division cycle, telomeres shorten by about 50-200 bp due to the end replication problem and active processing mechanisms (15–20). Cell arrest checkpoints are triggered once the telomeres reach a critically shortened length. With the ability of telomerase to maintain these ends, upon reactivation of telomerase the cells can continue to divide and acquire additional mutations in the process, eventually giving rise to a tumor (21). This is in line with findings where TPMs have been reported to be typically early events in cancers with high frequencies of these mutations (9, 22–25), behaving as a driver mutation in cancer via a two-step pathway (26). TPMs are initially gained in non-cancerous cells but do not participate actively in telomere extension. Once genome instability is triggered by multiple critically shortened telomeres, telomerase expression can be further upregulated subsequently in the presence of TPMs (26). *TERT* reactivation can also occur as a result of constitutive oncogenic cell signaling pathways, including TGF-β/Smad signaling and Wnt/β-Catenin signaling. Both can lead to upregulation of MYC, which binds E-boxes found in the *TERT* promoter to drive *TERT* expression (27–29). Likewise, Sp1 can bind to five Sp1 sites found in the *TERT* promoter and synergizes with MYC to increase *TERT* transcription (30, 31). These processes themselves are tightly regulated as illustrated by the TIP60-dependent acetylation of Sp1, which inhibits Sp1 binding to the TERT promoter and results in *TERT* repression (32). Furthermore, unlike conventional promoters, the majority of the CpG islands in the *TERT* promoter region are hypermethylated in cancer (33–35) except for a small non-methylated region upstream of the TSS, which coincides with the three-point mutations. Hypermethylation prevents the binding of transcriptional repressors such as CTCF and E2F1 while the non-methylated region grants access to activating transcription factors (34). The 11th, 12th, 19th and 27th CpG sites have been reported to be non-methylated in the active *TERT* promoter and are the binding sites for the activating transcription factors Sp1 (11, 12, 19) and MYC (27, 36). In comparison, non-cancerous primary cell lines are generally hypomethylated across the CpG islands (34, 37). *TERT* is also often expressed in a monoallelic manner, both in wildtype and TPM cell lines, with hypermethylation of promoter sequences being associated with repression and unmethylated alleles being expressed (37). In consequence, the expressed mutant allele in heterozygous TPM cell lines is associated with active histone marks such as H3K4me2/3, while the transcriptionally silenced wildtype allele is associated with repressive histone H3K27me3 marks (35, 38).

Most of the *TERT* promoter mutations result in the creation of a *de novo* site for ETS, a transcription factor family which consists of 29 genes (39). These proteins generally recognize the same GGA(A/T) motif, and they have been further categorized into twelve subgroups based on phylogenetic analysis of their DNA-binding domain (DBD) (40), with the ternary complex factor (TCF) subfamily recognizing a more specific binding motif of CCGGA(A/T) (41) that is also created by the C146T, C124T and A57C mutations. Bell *et al*. systematically tested 13 of the ETS family members that are expressed in glioblastoma multiforme (GBM) to determine which ETS member(s) was/were responsible for the upregulation of *TERT* expression in C146T and C124T-containing GBM cell lines (10). Here, GABPα (GA-binding protein α) was demonstrated to be the only critical member in the upregulation of the *TERT* gene specific to mutant C146T and C124T cell lines. GABP (GABPα+GABPβ) is the only obligate multimeric factor among the ETS family members (42) and is proposed to bind one of the two slightly overlapping endogenous ETS sites [ETS-96 and ETS-91 (**Figure 1A**)] alongside the *de novo* ETS site created by the C146T and C124T mutations. The same group would later demonstrate that knockdown of GABPβ1L, the long isoform of GABPβ1 (involved in GABP dimerization) also affected *TERT* expression in C146T and C124T GBM cell lines but not wildtype cell lines (43). Other factors that have been reported to regulate *TERT* expression in TPM dependent manner include p52 and ETS1/2 (C146T mutants only) (44) as well as phosphorylated ETS1 (due to constitutive MAPK pathway caused by NRAS/BRAF mutations in melanoma) (45).

**Figure 1.**
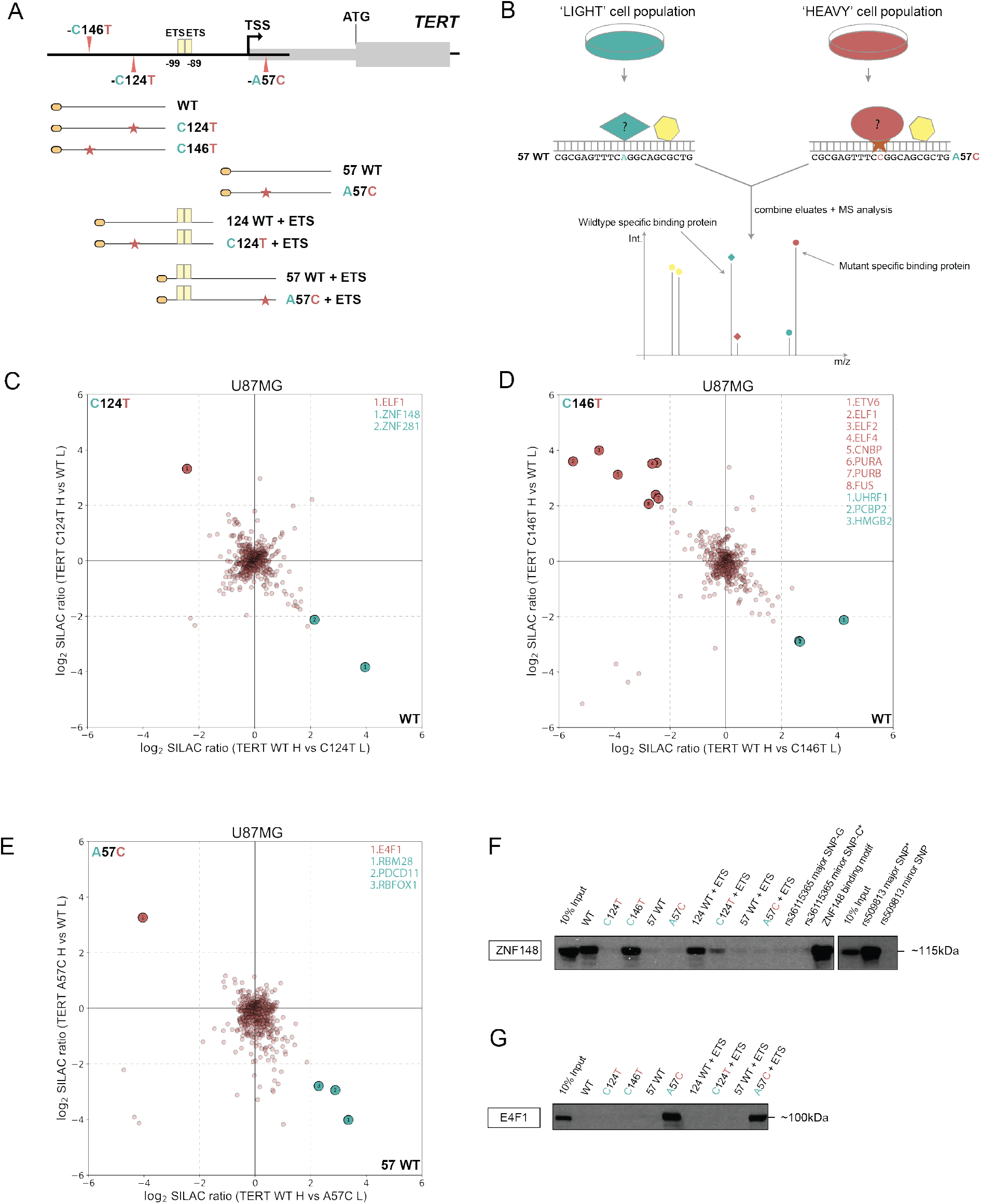
ZNF148 and E4F1 bind to wildtype and A57C *TERT* promoter sequence *in vitro*. **(A)** Schematic showing the coverage of the *TERT* promoter sequence by the DNA probes used in this study. **(B)** Workflow for a quantitative SILAC-based protein-DNA interaction screen. Biotinylated concatenated DNA oligonucleotides from panel A were immobilised on magnetic streptavidin beads and bound proteins were detected by MS or Western blot. 57 WT and A57C probes are shown as an example of wildtype and mutant sequences. Specific protein binders display a differential SILAC ratio, with background binders showing a 1:1 ratio. **(C)** Two-dimensional interaction plot for wildtype versus C124T using SILAC labelled U87MG nuclear protein extracts. Mutant-specific binders are labelled in red while wildtype-specific binders are labelled in green. **(D)** Two-dimensional interaction plot for wildtype versus C146T. Mutant- and wildtype-specific binders are labelled similar to (C). **(E)** Two-dimensional interaction plot for wildtype versus A57C. Mutant- and wildtype-specific binders are labelled similar to (C). **(F)** Sequence specific pull-down of endogenous ZNF148 with HeLa nuclear extracts using the 9 probes shown in (A) alongside rs36115365 major SNP-G and minor SNP-C, rs509813 major and minor SNP and CDKN1A/p21 promoter (containing ZNF148 binding site) probes. **(G)** Sequence-specific pull-down of endogenous E4F1 with HeLa nuclear extracts using the 9 probes shown in (A).

Although the aforementioned studies have managed to elucidate partly how TPMs can alter *TERT* transcription, we reasoned that a systematic screen would identify additional differential binders to wildtype and mutant *TERT* promoter and provide additional mechanistic insights underlying *TERT* expression control. Here, we performed *in vitro* DNA reconstitution pull-downs coupled with SILAC (stable isotope labeling with amino acids in cell culture)-based quantitative mass spectrometry analysis to identify differential binders between wildtype and the three mutant *TERT* promoter alleles (C146T, C124T and A57C). In this study, we demonstrated that ZNF148 binds preferentially to the wildtype *TERT* promoter sequence at position −124 and its binding is disrupted by the C124T mutation. Furthermore, for the A57C sequence we did not detect any ETS member but instead we identified E4F1 as an A57C mutant-specific transcription factor. Knockdown of both ZNF148 and E4F1 in corresponding wildtype and A57C mutant cell lines resulted in downregulation of *TERT* expression and concomitantly telomerase activity. Hence, ZNF148 and E4F1 are allele-specific *TERT* transcriptional activators, and may contribute to the deregulation of *TERT* expression in cancer.

## MATERIALS AND METHODS

### Cell culture and SILAC labeling

All cell lines (A375, HCT116, HeLa Kyoto, Saos2, U2OS, U87MG, 253J, T24, 575A, JON, HT1080 ST (46) and HEK293T) used in this study were cultured in 4.5 g/L glucose Dulbecco’s Modified Eagle Medium, supplemented with 10% FBS and 100 U/mL penicillin, 100 μg/mL streptomycin (Gibco). Cell lines were maintained at 37ºC with 5% CO_2_ in a humidified incubator.

For SILAC labelling, U87MG cells were cultured in DMEM (-Arg, -Lys) for SILAC (Thermo Scientific), supplemented with 10% dialyzed FBS (PAN-Biotech) and 100 U/mL penicillin, 100 μg/mL streptomycin (Gibco), in addition to either non-labelled 42 mg/L ^12^C_6_^14^N_4_-Arginine and 73 mg/L ^12^C_6_^14^N_2_-Lysine or heavy-labelled 42 mg/L ^13^C_6_^15^N^4^-Arginine and 73 mg/L ^13^C_6_^15^N_2_-Lysine. U87MG cells were cultured for at least two weeks in SILAC media prior to experiments to ensure an incorporation rate of >98%.

Ideal antibiotic concentrations for plasmid selection were optimized for each cell line by generating killing curves. The eventual concentrations used were 0.5 μg/mL for U87MG, 1 μg/mL for HeLa Kyoto, 575A, 2 μg/mL for T24 and 2.5 μg/mL puromycin for 253J. 10 μg/mL blasticidin S was used for 575A while 300 μg/mL hygromycin B were used for 575A.

### Sanger sequencing of cell lines

Genomic DNA (gDNA) was extracted from various cell line pellets using the QIAamp DNA Blood Mini Kit (Qiagen) according to the manufacturer’s instructions. The *TERT* promoter was then amplified with PCR using corresponding primers with the Q5 DNA polymerase (NEB) (**Supplementary Table S1**). PCR conditions were as follows: 98°C for 30 s, 35 cycles of 98°C for 10 s, 61°C for 25 s and 72°C for 40s. The PCR products were purified using the QIAquick PCR purification kit (Qiagen) according to the manufacturer’s protocol before being submitted for Sanger sequencing (AITBiotech). Chromatograms were inspected using 4Peaks (Nucleobytes).

### Plasmids and cloning

Full-length ZNF148, E4F1, ELF1, ELF2, GABPα and ZNF281 were amplified from HeLa Kyoto or U2OS cDNA (RNeasy plus mini kit; Qiagen) using the Q5 DNA polymerase (NEB) with specific primers (**Supplementary Table S1**). These sequences were cloned into the TOPO TA entry vector (Thermo Scientific) before LR recombination into Gateway-compatible expression vectors: pcDNA3.1 with N-terminal GFP tag or pcDNA-DEST47 with C-terminal GFP tag. Point mutants were generated using the Q5 site-directed mutagenesis kit (NEB) coupled with specific primers (**Supplementary Table S1**). Sequences for shRNA oligonucleotides were shortlisted from the GPP web portal (https://portals.broadinstitute.org/gpp/public/gene/search; BROAD institute) based on high specificity criteria. Primers (**Supplementary Table S2**) were annealed, phosphorylated and cloned into pLKO.1 vector following restriction enzyme digest and DNA ligation.

### Transfection and transduction

4-4.5 million HeLa Kyoto cells were seeded per 15 cm dish the evening prior to transfection for plasmid transfections. The cells were transfected with 121.6 μL of 1 mg/mL of linear Polyethylenimine (PEI; MW 25,000; Polysciences) and 30.4 μg of plasmid DNA diluted in Opti-MEM (Gibco). Media was replaced with fresh culture medium after 7 h and cells were harvested 48 h post-transfection.

For lentiviral production, 300,000 HEK293T were seeded per well in 6-well plate format the evening prior to transfection. Cells were transfected with 2.5 μL of 1mg/mL PEI, 0.5 μg of transfer vector (viral genome and gene of interest), 0.25 μg each of pMD2.G envelope plasmid (VSV-G), pRSV-Rev packaging plasmid (Rev) and pMDLg/pRRE packaging plasmid (Gag and Pol) diluted in Opti-MEM. Media was changed after 24 h and cell supernatants were harvested using a syringe and 0.45 μm filter unit 72 h post transfection.

For viral transduction, 100,000-200,000 cells were seeded per well in 6-well plates the evening prior to transduction. For every 1 mL of virus supernatant, 10 μL of 1 M HEPES (final 10 mM; Sigma-Aldrich) and 1 μL of 10 mg/mL (final 10 μg/mL) polybrene were added before addition to the cells to be transduced. These were replaced with fresh media 24 h post-transduction and selective media (puromycin, hygromycin or blasticidin) were added 48 h post transduction for cell selection.

### Reverse transcription and qPCR

Cells were harvested after 48 h (lentiviral transduction) + 72 h (puromycin selection). Cell pellets were washed with 1× PBS and RNA were extracted using the RNeasy plus mini kit (Qiagen) according to the manufacturer’s protocol. Total RNA was eluted in 30 μL nuclease-free water and used for cDNA conversion with the SuperScript IV Reverse Transcriptase (Thermo Scientific) based on the manufacturer’s protocol.

Each qPCR reaction consisted of 5 μL of 2× QuantiNOVA SYBR Green PCR master mix (Qiagen), 0.5 μL of 10 μM forward and reverse primers each, alongside appropriate cDNA amounts, topped up to 10 μL with nuclease-free water. Each reaction is set up in triplicates and ran on a QuantStudio3 or QuantStudio5 machine (Applied Biosystems), with the following protocol used: 2 min at 50°C, 10 min at 95°C, followed by 40 cycles at 95°C for 15 sec and 60°C for 1 min, then finally 95°C for 15 sec and 60°C for 1 min. mRNA levels of TBP were used as a housekeeping reference for the normalisation of mRNA levels of target genes (**Supplementary Table S3**).

### Nuclear protein extraction

Cells were harvested and washed in PBS and incubated on ice for 10 min with five times pellet volume of cold buffer A (10 mM HEPES pH 7.9, 10 mM KCl, 1.5 mM MgCl_2_). Cells were pelleted again and resuspended in approximately two times pellet volume of cold buffer A+ [buffer A supplemented with 0.1% IGEPAL CA-630, 0.5 mM DTT and 1× cOmplete protease inhibitor (Merck)] and homogenised in a glass dounce homogeniser (type B pestle) to mechanically lyse the cell cytoplasm. HeLa Kyoto cells required 40 dounces while 575A and U-87MG needed 30 dounces for maximum cytoplasmic lysis while keeping most nuclei intact. The supernatant (cytoplasmic fraction) was disposed as it was not required for this study. The nuclei were washed once with PBS and were then resuspended in 1.7× pellet volume of buffer C+ [20% (v/v) glycerol, 420 mM NaCl, 20 mM HEPES pH 7.9, 2 mM MgCl_2_, 0.2 mM EDTA pH 8, 0.1% IGEPAL CA-630, 0.5 mM DTT and 1× cOmplete protease inhibitor (Merck)] before being incubated on a rotating wheel at 30-35 rpm and 4ºC (cold room) for 1 h in order to extract nuclear proteins. The suspension was then centrifuged at maximum speed for 1 h at 4ºC to collect the nuclear extract. Protein quantification was performed with the Pierce BCA Protein Assay Kit (Thermo Scientific) according to the manufacturer’s protocol, and the nuclear protein extracts were diluted to the required concentrations using buffer C+.

### In vitro reconstitution DNA pull-down

DNA in vitro reconstitution pull-downs were essentially performed as previously described (47). 25 μL of 100 μM respective forward and reverse strand oligonucleotides (**Supplementary Table S4**) were denatured at 80ºC and re-annealed via gradual cooling. Double-stranded oligonucleotides were then concatemerized with 100 U T4 polynucleotide kinase (Thermo Scientific) and 20 Weiss U T4 DNA ligase (Thermo Scientific), biotinylated with desthiobiotin-ATP (Jena Bioscience) by 60 U Klenow fragment exo-(Thermo Scientific) and finally purified with Microspin G-50 columns (GE Healthcare). 0.25 mg (Western blot) or 0.75 mg (MS) Dynabeads MyOne Streptavidin C1 (Thermo Scientific) were washed twice with WB1000 buffer (1 M NaCl, 20 mM Tris, 0.1% IGEPAL CA-630 and 1 mM EDTA). The purified DNA probes were then immobilised on the streptavidin-coated magnetic beads for 30 min at RT on the rotating wheel. 100 μg (Western blot) or 300 μg (MS) of nuclear protein extracts alongside 20 μg (Western blot) or 40 μg (MS) of sheared salmon sperm DNA (Thermo Scientific), diluted in PBB buffer [(50 mM Tris,150 mM NaCl, 0.25% IGEPAL CA-630, 5 mM MgCl_2_ 1 mM DTT and 1× cOmplete protease inhibitor (Merck)] were added and incubated on the rotating wheel for 2 h at 4ºC. The beads were washed thrice with PBB, and bound protein were eluted in 2× Laemmli buffer and boiled for 5 min at 95ºC. Protein samples were separated on a 4-12% NuPAGE 4-12% Bis-Tris protein gels (Thermo Scientific) or 12% Bis-Tris protein gel (Thermo Scientific) for 1 h (Western blot) or 30 min (MS) at 170 V in 1× NuPAGE MOPS SDS running buffer (Thermo Scientific). The Colloidal blue staining kit (Thermo Scientific) was used to stain protein samples destined for downstream mass spectrometry analysis.

### Western blot

Cells were lysed in RIPA buffer supplemented with cOmplete protease inhibitor (Merck) by incubation on ice for 15 min. After centrifugation at 14,000 g, the supernatant was transferred to a fresh tube and quantified with Pierce BCA Protein Assay Kit (Thermo Scientific). 30-50 μg of protein extract were diluted in LDS sample buffer supplemented with 0.1 M DTT and boiled at 70ºC for 10 min. Protein samples were separated on a 4-12% NuPAGE 4-12% Bis-Tris protein gels (Thermo Scientific) for 1 h at 170 V in 1× NuPAGE MOPS SDS running buffer (Thermo Scientific). Separated proteins were transferred from the gel to a methanol-activated PVDF membrane for between 1-1.5 h at 70-220 mAmp. The membranes were blocked for 1 h at RT in PBS containing 5% (w/v) skim milk (Nacalai Tesque) and 0.1% Tween-20 (Nacalai Tesque) (PBS-T) and incubated with primary antibodies overnight at 4ºC or for 1 h at RT. Antibodies used are listed in **Supplementary Table S5**. Membranes were washed thrice in PBS-T and incubated with a secondary antibody for 1 h at RT. Chemiluminescence detection was performed using Pierce ECL Western blotting substrate (Thermo Scientific) or Amersham ECL prime Western blotting detection reagent (GE Healthcare).

### Mass spectrometry sample preparation, data acquisition and analysis

Following gel electrophoresis for 30 min, each lane was separated into four fractions. Gel pieces were then cut into approximately 1×1 mm pieces and destained twice with destaining buffer (50 mM NH_4_HCO_3_ and 50% ethanol), followed by dehydration with acetonitrile (ACN) and dried further in a speed vac. Reduction buffer (50 mM NH_4_HCO_3_, 10 mM DTT) was then added to rehydrate and reduce the samples at 56°C for 1 h. Reduction buffer was then replaced with alkylation buffer [50 mM NH_4_HCO_3_ and 55 mM ICH_2_CONH_2_ (Sigma-Aldrich)] and samples were alkylated in the dark for 45 min at RT. The gel pieces were then washed with ABC buffer (50 mM NH_4_HCO_3_), ACN, ABC buffer and ACN twice again, before drying further with a speed vac until the pieces were ‘dice-like bouncy.’ The protein samples were then incubated in digestion buffer [50 mM NH_4_HCO_3_ and 10 μg/mL Sequencing grade modified trypsin (Promega)] overnight at 37°C. Proteins were extracted and the supernatants were combined by incubating gel pieces in extraction buffer [35 mM ABC buffer, 30% ACN, 3% Trifluoroacetic acid], ACN, extraction buffer and ACN twice again. The combined supernatants were concentrated in a speed vac (Eppendorf) for about two hours before being transferred to pre-equilibrated stage tips. Stage tips were prepared by installing two C18 layers (Waters) inside a 200 μL pipette tip. The tips were washed with methanol and 80% ACN supplemented with 0.5% formic acid, then equilibrated by washing with 0.5% formic acid twice. After loading the combined supernatants, the tips were centrifuged followed by a wash with 0.5% formic acid. Tryptic peptides were eluted from stage tips with 80% ACN supplemented with 0.5% formic acid into a plate. Peptides were analysed by nanoflow liquid chromatography on an EASY-nLC 1200 system coupled to a Q Exactive HF mass spectrometer (Thermo Fisher Scientific). Peptides were separated on a C18-reversed phase column (25 cm long, 75 μm inner diameter) packed in-house with ReproSil-Pur C18-AQ 1.9 μm resin (Dr Maisch). The column was mounted on an Easy Flex Nano Source and temperature controlled by a column oven (Sonation) at 40°C. A 105-min gradient from 2 to 40% ACN in 0.5% formic acid at a flow of 225 nL/min was used. Spray voltage was set to 2.4 kV. The Q Exactive HF was operated with a TOP20 MS/MS spectra acquisition method per MS full scan. MS scans were conducted with 60,000 at a maximum injection time of 20 ms and MS/MS scans with 15,000 resolution at a maximum injection time of 50 ms. The raw files were processed with MaxQuant (48) version 1.5.2.8 with pre-set standard settings for SILAC labelled samples and the re-quantify option was activated. Carbamidomethylation was set as fixed modification while methionine oxidation and protein N-acetylation were considered as variable modifications. Search results were filtered with a false discovery rate of 0.01. Known contaminants, proteins groups only identified by site, and reverse hits of the MaxQuant results were removed and only proteins were kept that were quantified by SILAC ratios in both ‘forward’ and ‘reverse’ samples. The final SILAC plots were created with the aid of an in-house Python script. Heavy SILAC labelled input material was quality controlled to ensure a SILAC incorporation rate of >98%.

### Telomerase Repeat Amplification Protocol (TRAP)

Cells were lysed with two times the pellet volume of TRAP lysis buffer [1% IGEPAL CA-630, 50 mM Tris, 150 mM NaCl, 1× cOmplete protease inhibitor (Merck)] for 30 min on ice. The cells were then pelleted, and the supernatant was transferred to fresh tubes and quantified using the Pierce BCA Protein Assay Kit according to the manufacturer’s instructions. Each TRAP-qPCR was set up in a 20 μL reaction, consisting of 10 μl of 2× QuantiNOVA SYBR Green PCR master mix (Qiagen), 7.2 μL nuclease-free water, 0.4 μL of 10 μM (μmol/L) TS primer and ACX primer each and 2 μL of 250 ng/μL or 25 ng/μL protein sample (500 ng or 50 ng total nuclear extract respectively). All reactions were done in triplicates. The PCR protocol was as follows: 30 min at 25°C for reverse transcription by telomerase, then heat inactivation and hot-start of Taq polymerase at 95°C for 2 min, followed by 32 cycles of 95°C for 5 sec (denaturation) and 60°C for 90 sec (annealing and elongation), and finally held at 10°C. Cell lysates from telomerase-negative cells (U2OS and Saos2) and heat-inactivated lysates (85°C for 10 min) were used as negative controls while HT-1080 ST lysates were used as positive controls. qPCR analysis was performed using Thermo Scientific software.

For TRAP-ddPCR (digital droplet PCR), the telomerase extension was performed first, with each reaction being set up with 5 μL of 10× TRAP extension buffer [630 mM KCl, 200 mM Tris, 15 mM MgCl_2_, 0.5% (v/v) TWEEN 20 and 10 mM EGTA], 1 μL of 10 μM TS primer, 1 μL of 10 mM dNTP mix, 500 ng of nuclear protein sample topped up to 40 μL with nuclease-free water. The samples were incubated at 25°C for 30 min, 95°C at 5 min and cooled to 4°C. To set up the ddPCR, each reaction consists of 11 μL 2× QX200 ddPCR EvaGreen Supermix, 0.11 μL of 10 μM TS primer and ACX primer each were added to 2.2 μL of telomerase-extended products topped up to 22 μL with nuclease-free water. Oil droplets were generated using the Automated Droplet Generator (Bio-Rad), followed by plate sealing with a pierceable foil heat seal. A PCR was conducted with the following conditions, 95°C for 5 min, followed by 40 cycles of 95°C for 30 sec, 54°C for 30 sec and 72°C for 30 sec, and finally held at 4°C with a ramp-rate of 2.5°C/sec between each step. Finally, the plate was transferred to a QX200 Droplet Reader (Bio-Rad) and positive droplets were analyzed using the QuantaSoft software (Bio-Rad).

### Pyrosequencing

For pyrosequencing, 500 ng of genomic DNA was subjected to bisulfite conversion using the EpiTect Bisulfite Kit (Qiagen). PCR reactions were performed on 12.5 ng of bisulfite-treated DNA in a final volume of 25 μL using the Pyromark PCR kit (Qiagen) with one of the primers being biotinylated for later capture. The primers were designed using the PyroMark Assay Design Software 2.0 (Qiagen) (**Supplementary Table S6**). The initial denaturation/activation step was performed at 95°C for 15 min, followed by 50 cycles of 30 sec at 94°C, 30 sec at 54°C, 45 sec at 72°C and a final extension step at 72°C for 10 min. The quality and the size of the PCR products were evaluated by running 5μL of each PCR product on 1.5% (w/v) agarose gel in a 0.5X TBE buffer. The biotinylated PCR products were immobilized on streptavidin-coated sepharose beads (GE Healthcare). DNA strands were separated using the PyroMark Q24 Vacuum Workstation, the biotinylated single strands were annealed with 0.375 μM sequencing primer (**Supplementary Table S6**) and used as a template for pyrosequencing. Pyrosequencing was performed using PyroMark Q24 Advanced (Qiagen) according to the manufacturer’s instructions, and data about methylation at each CpG was extracted and analyzed using the PyroMark Q24 Advanced 3.0.0 software (Qiagen).

## RESULTS

### ZNF148 and E4F1 bind to the wildtype and A57C *TERT* promoter *in vitro*

In order to identify factors that are binding to either the wildtype (WT) or mutant (C146T, C124T and A57C) *TERT* promoter, we designed different DNA probes, each containing either the wildtype or mutant *TERT* promoter sequence (**Figure 1A**; refer to **Supplementary Table S4** for exact sequences). We then performed *in vitro* DNA reconstitution pull-downs with the wildtype and mutant probes combined with SILAC-based quantitative mass spectrometry analysis (**Figure 1B**) with either light-labelled or heavy-labelled U-87MG (containing the C124T mutation; **Supplementary Table S7**) nuclear protein extracts. Corresponding eluates were combined and analysed using quantitative mass spectrometry to identify proteins that were binding preferentially to either the wildtype (bottom right quadrant) or mutant (top left quadrant) *TERT* promoter sequences (**Figures 1C-E**). Only proteins specifically enriched with SILAC ratios >4 in both the ‘forward’ and ‘reverse’ experiments were considered to be positive hits (**Figures 1C-E; Supplementary Tables S8-10**).

Consistent with previous reports (1, 2) that predicted the creation of a *de novo* ETS motif, several ETS factors were identified binding preferentially to C124T (ELF1) and C146T (ELF1, ELF2, ELF4, ETV6) mutant sequences (**Figures 1C, D; Supplementary Tables S8-9**). GABPα was not identified, consistent with a similar screen that required the introduction of a second ETS site to observe GABPα binding (not included in our probes used for mass spectrometry analysis) (49). Although the A57C mutation also creates an ETS binding motif, no ETS factors were identified binding preferentially to the mutant sequence. Instead we identified E4F1 as the only A57C-specific candidate protein (**Figure 1E; Supplementary Table S10**). The design of our experiment allowed us to read out both mutant- and wildtype-specific binding proteins simultaneously. In agreement with a recent report (50) ZNF148 and ZNF281 were specifically enriched on the WT probe in comparison to the C124T mutant and UHRF1 on the WT probe compared to C146T (**Figures 1C, D; Supplementary Tables S8-9**). Notably, some known single-stranded DNA/RNA binding proteins (including CNBP, PURA, PURB, FUS, PCBP2, HMGB2, RBM28, PDCD11 and RBFOX1) were also identified to preferentially bind to either wildtype or mutant sequence (**Figures 1D, E; Supplementary Tables S9-10**), which may be due to the presence of some single-stranded DNA as part of our concatenated DNA probe preparation. We next focused on proteins with a DNA binding domain that are more likely to bind directly to the dsDNA sequences.

In order to systematically validate the absolute enrichment of all wildtype or mutant specific binders, HeLa (wildtype *TERT* promoter; **Supplementary Table S7**) nuclear protein extracts were incubated with the five probes used above, alongside WT124+ETS/C124T+ETS and WT57+ETS/A57C+ETS (**Figure 1A**), followed by Western blot. The additional probes contain the two endogenous ETS sites between −99 and −89. While a previous report (49) using a similar pull-down approach had reported GABPα binding in the presence of both the endogenous ETS sites and *de novo* ETS site created by the C124T mutation using UACC903 (melanoma; C124T mutant) nuclear extracts (**Figure 1A, Supplementary Table S4**), we could not detect GABPα enrichment to any of the nine probes using antibodies against endogenous GABPα, consistent with our mass spectrometry results. Similar results were obtained for pull-downs using A375 (melanoma, C146T mutant; **Supplementary Table S7**) nuclear extracts or with HeLa cells transiently transfected with both N-terminally and C-terminally tagged GFP-GABPα. In all cases, we were unable to enrich for GABPα (**Supplementary Figure 1A**), which might be partly attributed to cell-type-specific effects. In contrast, N-terminally tagged GFP-ELF1 showed strong absolute enrichment on the C146T probe, while N-terminally tagged GFP-ELF2 showed strong absolute enrichment on all mutant probes as compared to their respective wildtype (**Supplementary Figure 1A**), confirming the idea that ETS family members may represent the predominant transducers of C124T- and C146T-dependent *TERT* expression. The fact that they were enriched both in our mass spectrometry and Western blot experiments in the absence of the additional endogenous ETS sites suggests that their binding, at least *in vitro*, only requires the TPM alleles.

Furthermore, both endogenous ZNF148 and N-terminally tagged GFP-ZNF148 were enriched on the WT sequence (WT and WT124+ETS) but did not bind to the C124T and C124T+ETS mutant probes (**Figures 1F, 2B**). ZNF148 has been reported to be an allele-specific transcription factor in two previous studies (51, 52), shown to preferentially bind one single nucleotide polymorphism (SNP) allele over the other. We decided to include both SNPs in our probes design (rs509813 and rs36115365) alongside a known ZNF148 binding site (p21 promoter) (**Supplementary Table S4**) (51). Differential ZNF148 binding to the rs36115365 SNP was particularly intriguing since it is located 18 kb upstream of the 5’- end of *TERT* and its knockdown in a panel of pancreatic cancer cell lines resulted in reduced *TERT* expression. However, we could not recapitulate binding to the rs36115365 SNP using HeLa nuclear extracts, while we could readily see differential binding of ZNF148 to the rs509813 major SNP and the p21 promoter binding site in the same pulldown assay (**Figures 1F, 2B**). While the lack of ZNF148 binding to the rs36115365 may again in part be attributed to cell-type-specific differences, the identification of ZNF148 binding to the proximal promoter sequence might implicate it in direct *TERT* mRNA expression regulation independent of long-ranged chromatin-chromatin interactions. Although ZNF281 displayed a similar binding profile as ZNF148 (**Supplementary Figure 1A**), the absolute enrichment of ZNF281 on the wildtype probes were much weaker as compared to 10% input, which might imply a low overall affinity to the TERT promoter sequence. ZNF281 was hence excluded from functional downstream analyses.

Finally, we could validate E4F1 binding to the A57C mutant sequence using both an antibody against endogenous E4F1 and C-terminally tagged E4F1-GFP on both the A57C and A57C+ETS probes as compared to their respective wildtype controls with strong absolute enrichment (**Figures 1G, 2C**). These data collectively demonstrate that ZNF148 and E4F1 specifically bind to the wildtype and A57C *TERT* promoter sequences *in vitro*, respectively.

### ZNF148 and E4F1 bind directly to WT and A57C *TERT* promoter respectively

To test whether ZNF148 and E4F1 bind directly to the respective *TERT* promoter sequences, DNA binding mutant constructs of both ZNF148 and E4F1 were generated by mutating key cysteine residues in the C2H2 zinc finger motifs in both proteins and were then tested in our DNA pull-down assay (53) **(Figures 2A-C)**. Mutation in either of the four zinc fingers or deletion of all four zinc fingers in ZNF148 resulted in the loss of ZNF148 binding to the WT, C146T, 124WT+ETS and ZNF148 binding motif probes (**Figure 2B**). Similarly, mutation in either of the first two zinc fingers in E4F1, previously shown to be critical for its DNA binding ability (54), resulted in the loss of E4F1 binding to WT57 and WT57+ETS probes (**Figure 2C**). These results suggest that both factors likely bind the *TERT* promoter directly. We also aligned the respective *TERT* promoter alleles to the binding motifs of ZNF148 and E4F1 as predicted by the MethMotif database (55) and identified a high degree of overlap between the general motifs and the exact binding sites in the *TERT* promoter. Importantly, the mutations overlap with key residues, in which alteration of one base results in the disruption (C124T) and creation (A57C) of the ZNF148 and E4F1 binding motif, respectively **(Figure 2D)**. These data further strengthen a model in which ZNF148 and E4F1 bind directly to the WT and A57C mutant *TERT* promoter sequences.

**Figure 2.**
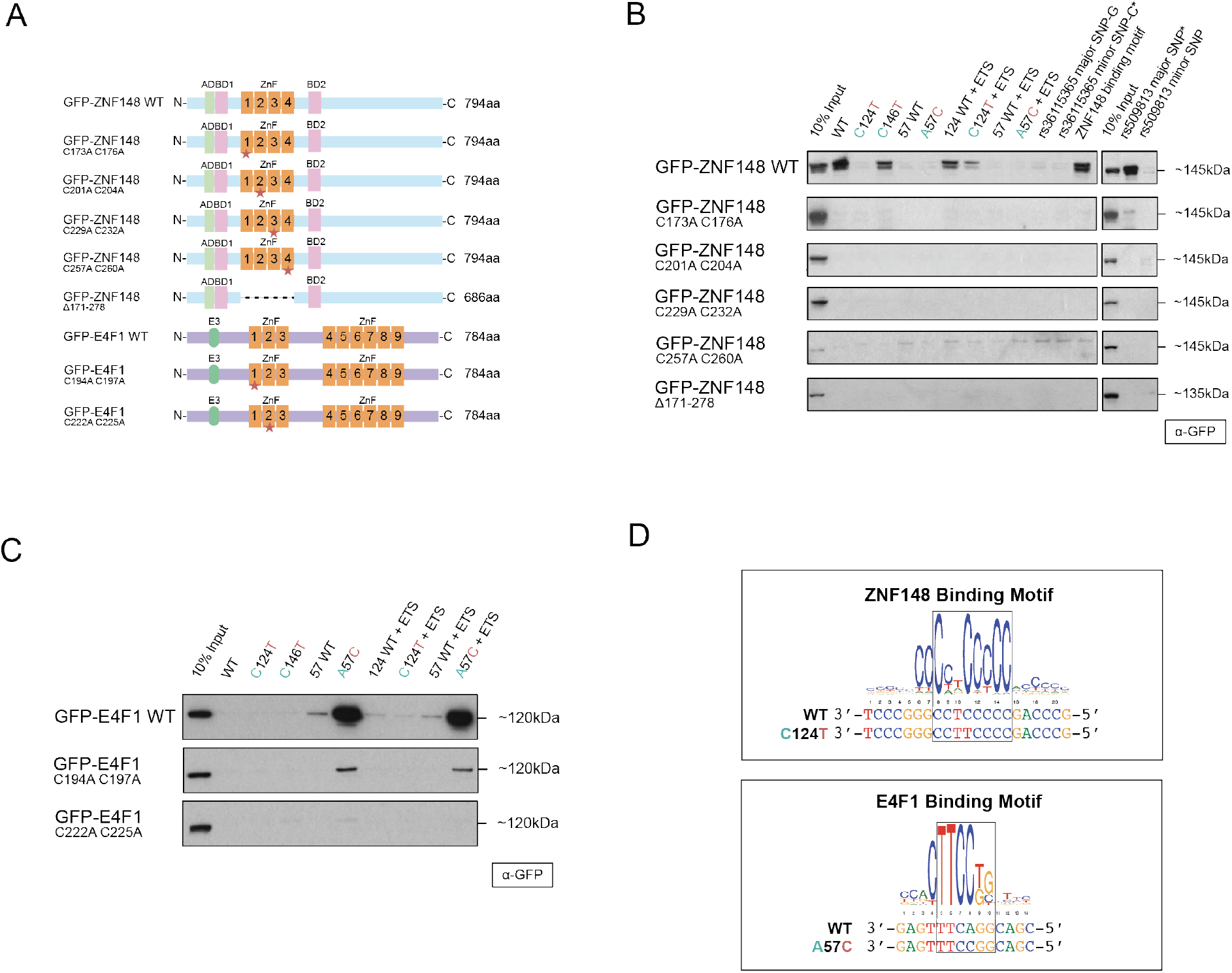
ZNF148 and E4F1 require their DNA binding domains to bind *in vitro* to WT and A57C *TERT* promoter, respectively. **(A)** Schematic representation of ZNF148 and E4F1 WT and DNA binding mutant constructs. **(B)** Sequence-specific pull-down of recombinant GFP-tagged wildtype or binding mutant of ZNF148 with HeLa nuclear extracts with the 9 probes shown in Fig. 1A alongside rs36115365 major SNP-G and minor SNP-C, rs509813 major SNP and minor SNP and CDKN1A/p21 promoter (containing ZNF148 binding site). **(C)** Sequence-specific pull-down of recombinant GFP-tagged wildtype or binding mutant of E4F1 with HeLa nuclear extracts with the 9 probes shown in Fig. 1A. **(D)** Binding motif of ZNF148 and E4F1 aligned to the *TERT* promoter sequence based on the consensus motif derived from the MethMotif database (55).

### ZNF148 and E4F1 do not affect TERT promoter methylation

To study whether the DNA methylation pattern is altered between cell lines with different TPM status, pyrosequencing was conducted using gDNA extracted from the different cell lines. The DNA methylation pattern at the CpG islands across the *TERT* promoter is different dependent on the mutation status of the cell lines. In general, WT cell lines (HeLa Kyoto and 253J, excluding HCT116 which displays hypomethylation) display a higher degree of methylation as compared to cell lines with either C124T (T24 and U87MG) or C146T (A375) mutation (**Supplementary Figure 2**). The two cell lines with A57C mutation displayed more variation in their DNA methylation levels, with JON showing low DNA methylation pattern across its *TERT* promoter, that is closer to IMR90 (immortalized normal fibroblasts, where normal cells typically display hypomethylation across the *TERT* promoter) while 575A is closer to WT cell lines. Of note, we observed a drop in DNA methylation levels especially around the region where both C124T and C146T mutation reside (between CG7 to CG10; which includes the third Sp1 site and near the second and fourth Sp1 sites) in all cell lines despite their mutation statuses. In comparison, the decrease in DNA methylation levels is not as pronounced for the A57C mutation locus (between CG17 and CG18) across all cell lines. These data may indicate generally greater accessibility at the key binding sites for TPM-specific transcription factors (1, 2, 35, 38).

To test if ZNF148 and E4F1 might affect the epigenetics of the *TERT* promoter, pyrosequencing was performed to study the effects on DNA methylation following shRNA knockdown. No significant differences were observed in DNA methylation of the proximal *TERT* promoter following shRNA knockdown of ZNF148 in HeLa Kyoto and E4F1 in 575A (**Figure 3**). In addition, no effects on DNA methylation were observed following the knockdown of other known transcription factors of the *TERT* promoter (MYC, SP1, GABPα, GABPβ1L) in the two cell lines **(Figure 3)**, suggesting that these factors do not act on the TERT promoter by modulating the DNA methylation status.

**Figure 3.**
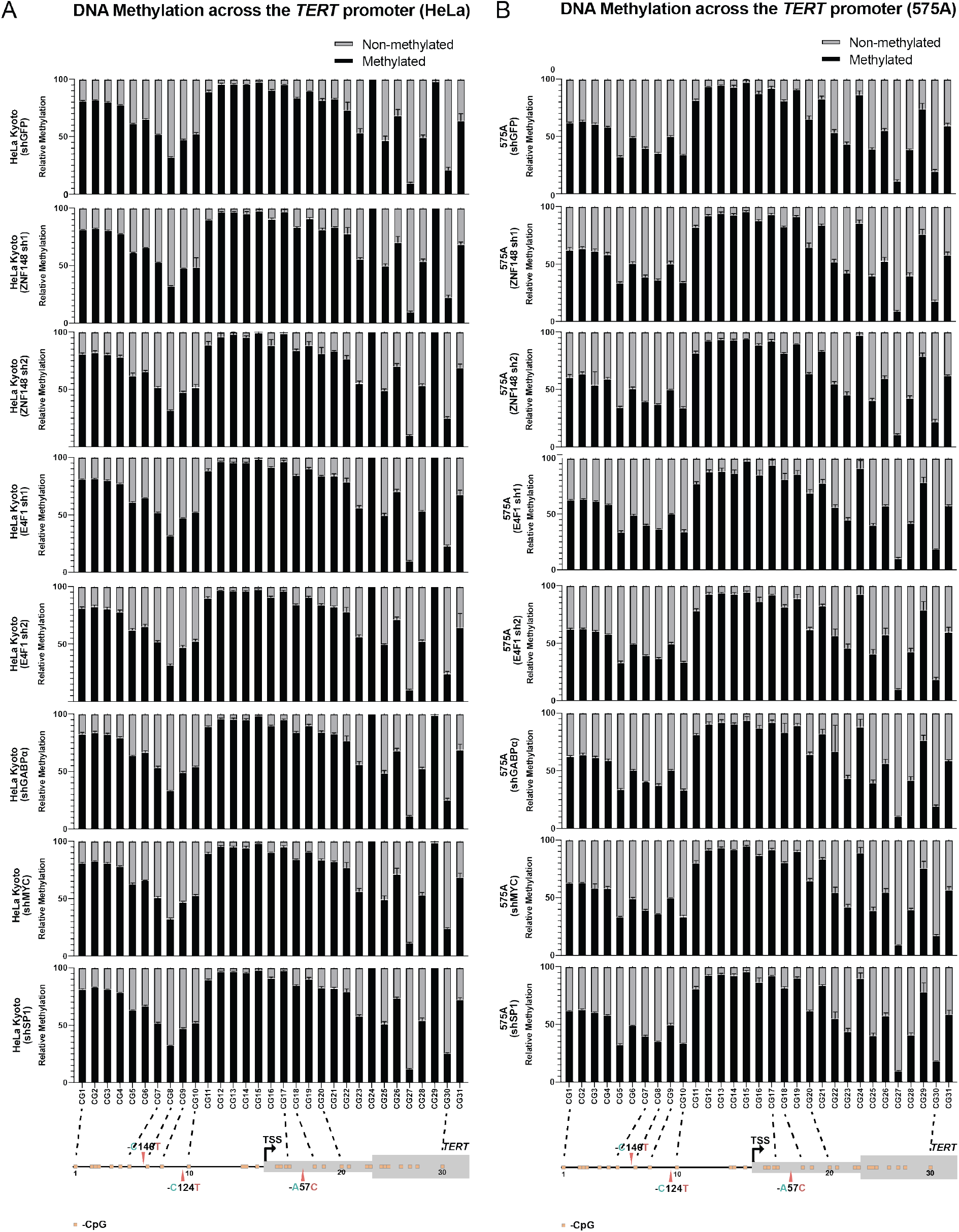
TERT promoter mutant-specific binders do not affect the promoter methylation status. **(A)** Relative DNA methylation frequency of CpGs across the *TERT* promoter in HeLa Kyoto cells following shRNA knockdown. **(B)** Relative DNA methylation frequency of CpGs across the *TERT* promoter in 575A cells following shRNA knockdown.

### ZNF148 and E4F1 act as transcriptional activators of *TERT*

To test if the promoter binding of ZNF148 and E4F1 translates into an actual effect on *TERT* transcript levels, we depleted both factors with two shRNAs each (**Figures 4, 5; Supplementary Figure 3**) across a panel of cell lines with either wildtype or mutant *TERT* promoters (**Supplementary Table S7**). In addition, we used shRNAs targeting TERT, GABPα, GABPβ1L, MYC and Sp1 as positive controls. MYC and Sp1 are factors that have previously been reported to bind to the E-box and Sp1 sites on the *TERT* promoter (56–58) and affect *TERT* transcription while knockdown of GABPα and GABPβ1L have been reported to only affect *TERT* transcription in cell lines containing the C124T or C146T mutation (10, 43). Upon ZN148 knock-down in HeLa cells, we observed a reduction of *TERT* expression for both shRNAs (**Figure 4A**). As expected, shTERT, shMYC and shSp1 also resulted in reduction of *TERT* expression. Albeit HeLa cells being TERT promoter WT, knockdown of GABPα also led to a statistically significant reduction of *TERT*. It is of note that knockdown of all other factors excluding ZNF148 resulted in an increase in ZNF148 mRNA and protein levels (**Figures 4B, 5B, Supplementary Figure 3A**), suggesting some degree of interdependency or feedback regulation. Since we could not identify a commercially available TERT antibody that showed depletion of a specific band upon shTERT transduction, we used the Telomerase Repeat Amplification Protocol (TRAP) assay as an indirect measure for changes in TERT protein levels. HT1080 Super Telomerase cells (HT1080 ST) were used as a positive control while the telomerase-negative cell lines U2OS and Saos2 were used as negative controls alongside heat-inactivated cell extracts. Using HeLa Kyoto cells, we observed a reduction of telomerase activity for both shRNAs targeting ZNF148 whereas there was a marginal effect caused by shGABPα and shGABPβ1L, consistent with previous reports where these factors only affect cell lines with C146T or C124T mutations. Similarly, we also saw a reduction in telomerase activity with our positive controls shTERT, shMYC and shSp1 (36, 59–61) (**Figure 4C**). We also observed a similar trend of reduction in *TERT* mRNA and telomerase activity in another wildtype *TERT* promoter cell line, 253J (**Supplementary Table S7**), following ZNF148 shRNA knockdown (**Supplementary Figures S3B-C**), concomitant with a reduction in *TERT* expression and telomerase activity upon TERT, MYC and Sp1 knockdown. Again, both GABPα and GABPβ1L knockdown had marginal effects on *TERT* mRNA expression, although shGABPβ1L led to a more pronounced reduction in telomerase activity. In contrast, ZNF148 knockdown in C124T-positive T24 cells (**Supplementary Table S7**) had marginal effects on *TERT* mRNA expression and telomerase activity, in agreement with a lack of promoter binding in the presence of C124T mutations (**Supplementary Figures S4D-E**) while our positive controls shTERT and shMYC both resulted in a consistent reduction of *TERT* mRNA expression and telomerase activity.

**Figure 4.**
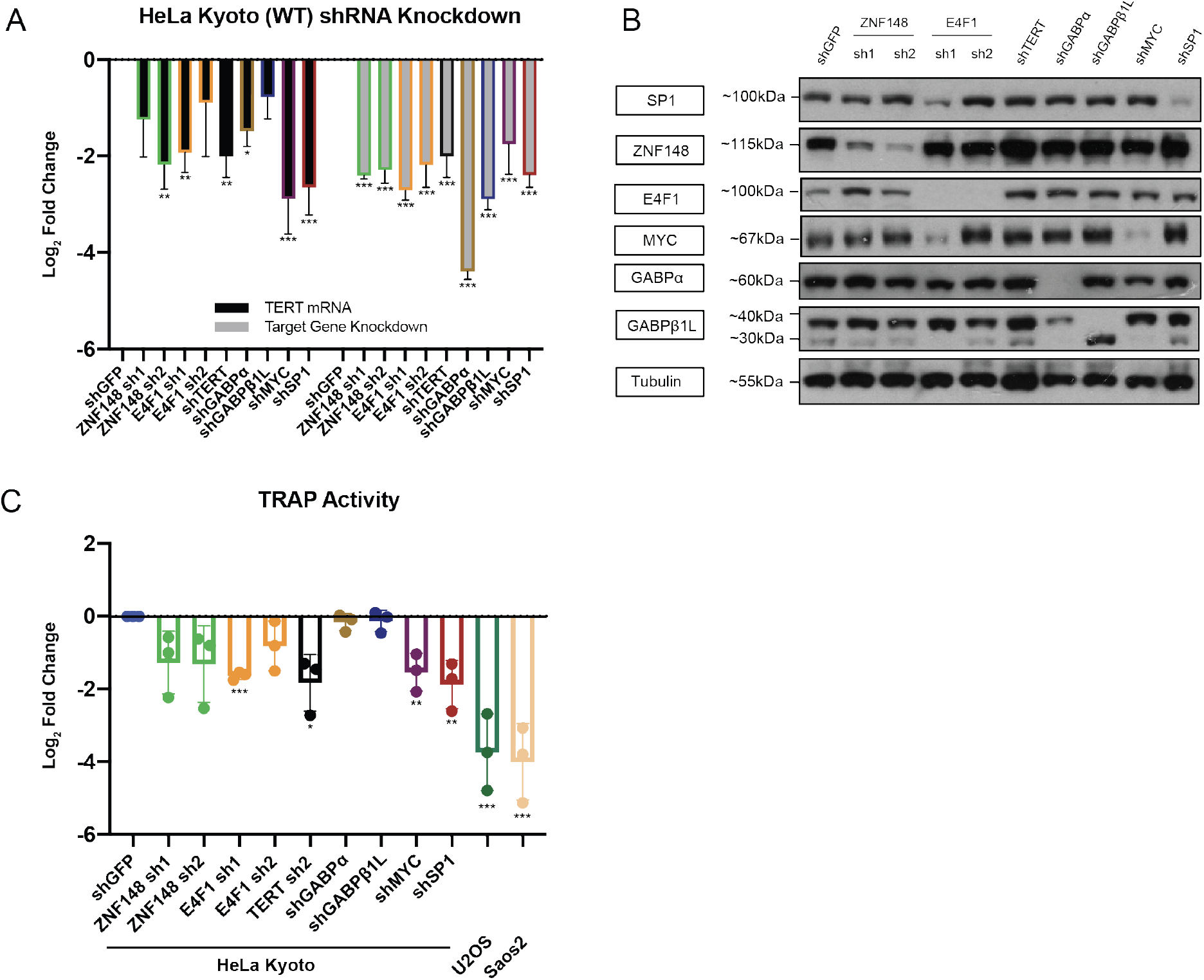
ZNF148 knockdown led to reduction of *TERT* promoter transcription and telomerase activity in HeLa cells. **(A)** mRNA expression data of *TERT* and target genes following 48+72 h (48h virus transduction, 72 hours puromycin selection) post-shRNA knockdown in HeLa, with shGFP as control. Data shown as mean of values from three biological replicates. **(B)** Representative Western blot image following 48+72 h post-shRNA knockdown in HeLa, with shGFP as control. **(C)** TRAP assay measuring telomerase activity following 48+72 h post-shRNA knockdown in HeLa, with shGFP as control. Telomerase-negative U2OS and Saos2 were used as negative controls. Data shown as mean of values from three biological replicates. All statistical significance was calculated using a two-sampled t-test, and the degree of significance is indicated as: * for p <0.05; ** for p <0.01; *** for p <0.001.

**Figure 5.**
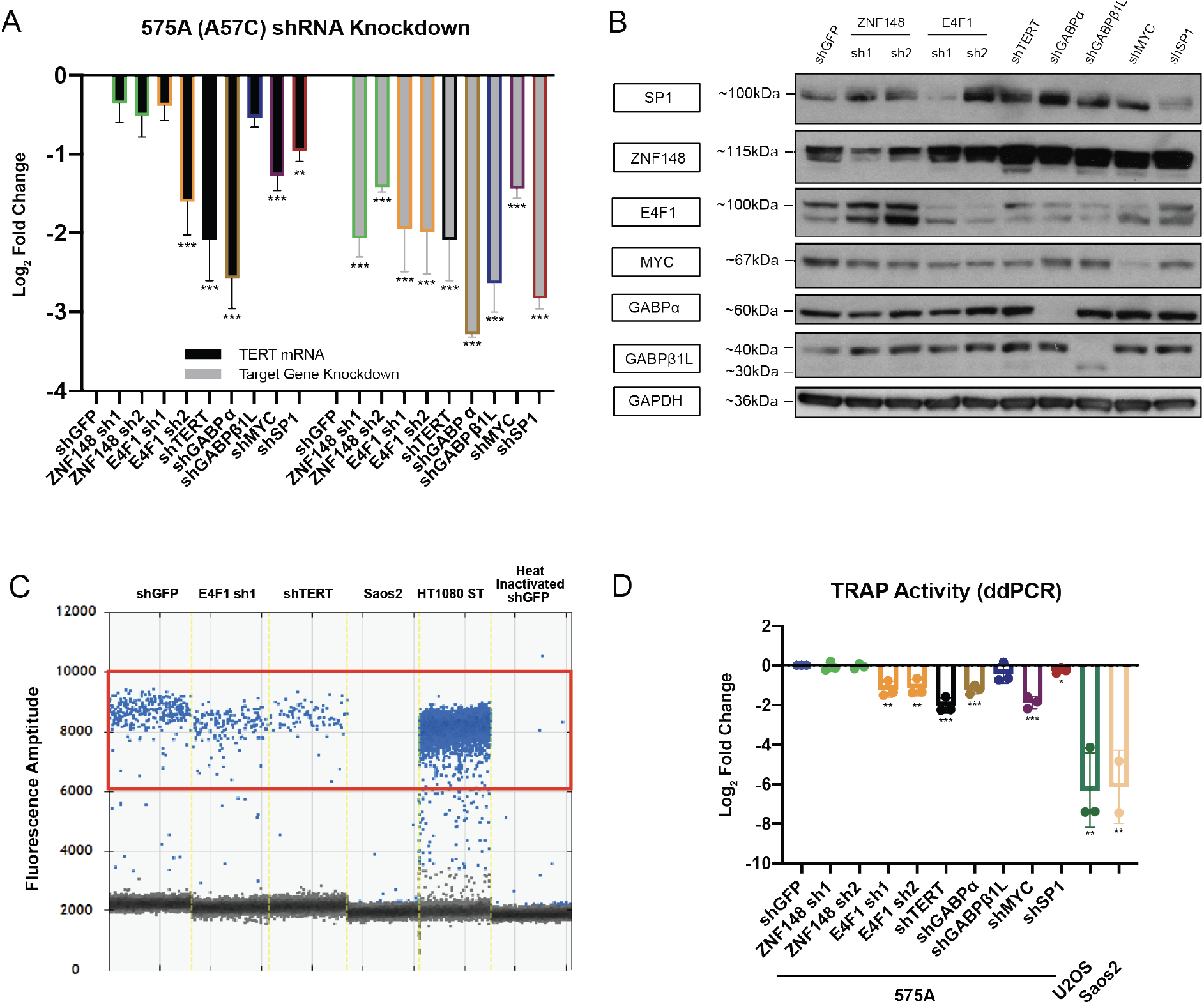
E4F1 knockdown led to reduction of *TERT* promoter transcription and telomerase activity in 575A cells. **(A)** mRNA expression data of *TERT* and target genes following 48+72 h (48h virus transduction, 72 hours puromycin selection) post-shRNA knockdown in 575A, with shGFP as control. Data shown as mean of values from three biological replicates. **(B)** Representative Western blot image following 48+72 h post-shRNA knockdown in 575A, with shGFP as control. **(C)** ddPCR-TRAP assay measuring telomerase activity following 48+72 h post-shRNA knockdown in 575A, with shGFP as control. Telomerase-negative U2OS and Saos2 were used as negative controls. Positive droplets quantified are labeled in blue. **(D)** Quantification of data in (C), data shown as mean of values from three biological replicates. All statistical significance was calculated using a two-sampled t-test, and the degree of significance is indicated as: * for p <0.05; ** for p <0.01; *** for p <0.001.

In A57C-positive 575A cells (**Supplementary Table S7**), in addition to shTERT and shMYC, knockdown of E4F1 resulted in a general reduction of *TERT* mRNA expression (**Figure 5A, B**) and TRAP activity as measured by digital droplet PCR (ddPCR) (62, 63). The latter showed a clear reduction of positive droplets for shE4F1 and shTERT as compared to shGFP (control), with an abundance of droplets in HT1080 ST and absence of droplets for U2OS and Saos2 alongside heat-inactivated shGFP (**Figures 5C, D**). Therefore, the knockdown of ZNF148 in a wildtype *TERT* promoter cell line (HeLa Kyoto) and the knockdown of E4F1 in the presence of the A57C mutation resulted in reduction of telomerase activity. These data demonstrate a functional effect of these transcription factors at their respective *TERT* promoter loci and are in line with allele-specific regulation.

## DISCUSSION

We demonstrated that a SILAC-based DNA-protein interaction assay could be a highly viable and robust method for the identification of novel binding factors to wildtype and mutant sequences. The binding of ETS factors such as ELF1 and ELF2 to the *de novo* ETS site created by the mutant sequences (**Supplementary Figure 1A**) here serves as a proof-of-concept. Although all three TPMs create the same basic ETS consensus site, no ETS factors were enriched on the mutant A57C *TERT* promoter (**Figure 1E**) in our proteomics screen. This may be due to the differences in flanking sequences between the A57C mutant (gccGGAAactc) and the C124T and C146T mutants (cccGGAAgggg) despite the common GGAA motif, in agreement with previous reports demonstrating the importance of flanking sequences in determining which ETS family members are recruited to specific loci (64).

Next, we were unable to reproduce the enrichment of GABPα using probes that contain the endogenous ETS sites alongside the *de novo* C124T mutation as previously reported (49). We were also unable to recapitulate the enrichment of ZNF148 on the minor rs36115365 SNP using the exact same pulldown assay as a previous report (51), which might be partly attributed to different wash conditions (e.g. buffer salt concentration) and incubation duration as well as cell line specific effects. In the latter study, the authors demonstrated that knockdown of ZNF148 led to a reduction of *TERT* transcription in both pancreatic cancer cell lines containing the minor SNP and those with the major SNP. This lends support to our work, implying ZNF148 as a direct transcriptional activator at the proximal wildtype *TERT* promoter. If ZNF148 binding to the rs36115365 is context-specific, binding to both the distal variant and the proximal promoter could lead to a ZNF148-mediated chromatin-chromatin interaction to further strengthen the transcriptional output.

In this study, we have identified an A57C TPM-specific transcriptional activator: E4F1 has been previously reported to be involved in carcinogenesis by regulating viral oncoprotein expression (65, 66), p53 ubiquitination (53) as well as interacting with several tumor suppressors (p14ARF, BMI1, pRB and RASSF1A) and oncogenes (HNF1, HMGA2, Smad4 and TCF3) (67). It also directly controls the transcription of multiple mitochondrial and checkpoint protein genes, including DNA-damage response protein Chek1 (67). The oncogenic behaviour is consistent with our study, in which the A57C TPM creates a *de novo* binding site for E4F1 to upregulate *TERT* oncogene expression.

Based on our knockdown experiments, we postulate that there might be possible crosstalk between the different transcription factors that regulate *TERT* expression. For instance, the upregulation of ZNF148 following knockdown of all other factors, including oncoprotein E4F1, could be a compensatory mechanism to upregulate *TERT* expression (**Figure 4B, 5B**), which is vital for unlimited cancer cell proliferation.

In conclusion, we have identified two novel transcriptional regulators of the *TERT* locus, which contribute to *TERT* upregulation in a mutation-dependent manner. Specifically, we could identify ZNF148 as an additional factor that could regulate the *TERT* promoter at the −124 position, with corresponding decrease in *TERT* expression and telomerase activity upon ZNF148 knockdown in wildtype *TERT* promoter cell lines. Additionally, we here report the first transcription factor that specifically regulates *TERT* expression at the A57C mutant promoter. Our study alongside others (49, 51, 68, 69) demonstrates the potential of quantitative mass spectrometry in combination with *in vitro* reconstitution as a systematic strategy to interpret non-coding mutations.

## DATA AVAILABILITY

The mass spectrometry proteomics data have been deposited to the ProteomeXchange Consortium via the PRIDE partner repository (70) with the dataset identifier PXD037776.

## ACKNOWLEDGMENTS

We are grateful to all members of the Kappei lab for advice and discussions. 253J, T24, 575A and JON were kindly provided by Dr. Kees Jansen, Radboud University Medical Centre, Nijmegen, Netherlands.

## AUTHOR CONTRIBUTIONS

BHC and DK conceived the study and designed experiments. BHC performed experiments with help from LF, CD, NZA, AW and FB. DGT, SJ and PAD contributed to the research supervision. BHC and DK analysed the data. BHC and DK wrote the manuscript with input from all authors.

---

**Supplementary Figure S1.**
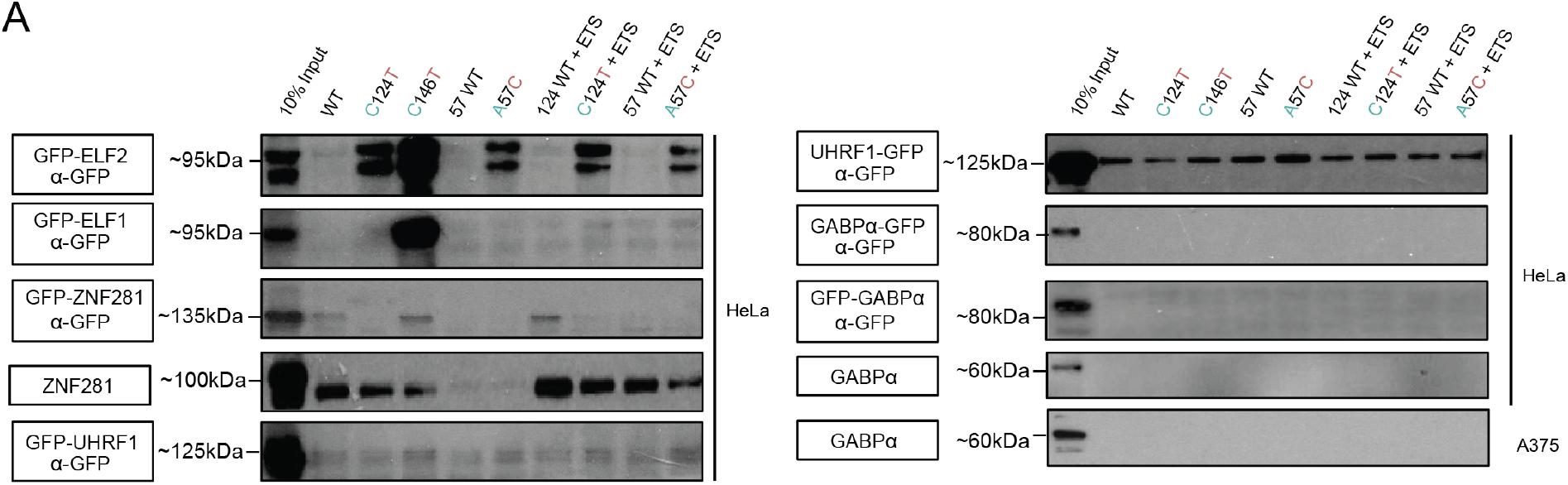
**(A)** Sequence specific pull-down of endogenous and/or recombinant GFP-tagged ELF1, ELF2, ZNF281, UHRF1 and GABPα with HeLa and/or A375 nuclear extracts using the 9 probes shown in Fig. 1A.

**Supplementary Figure S2.**
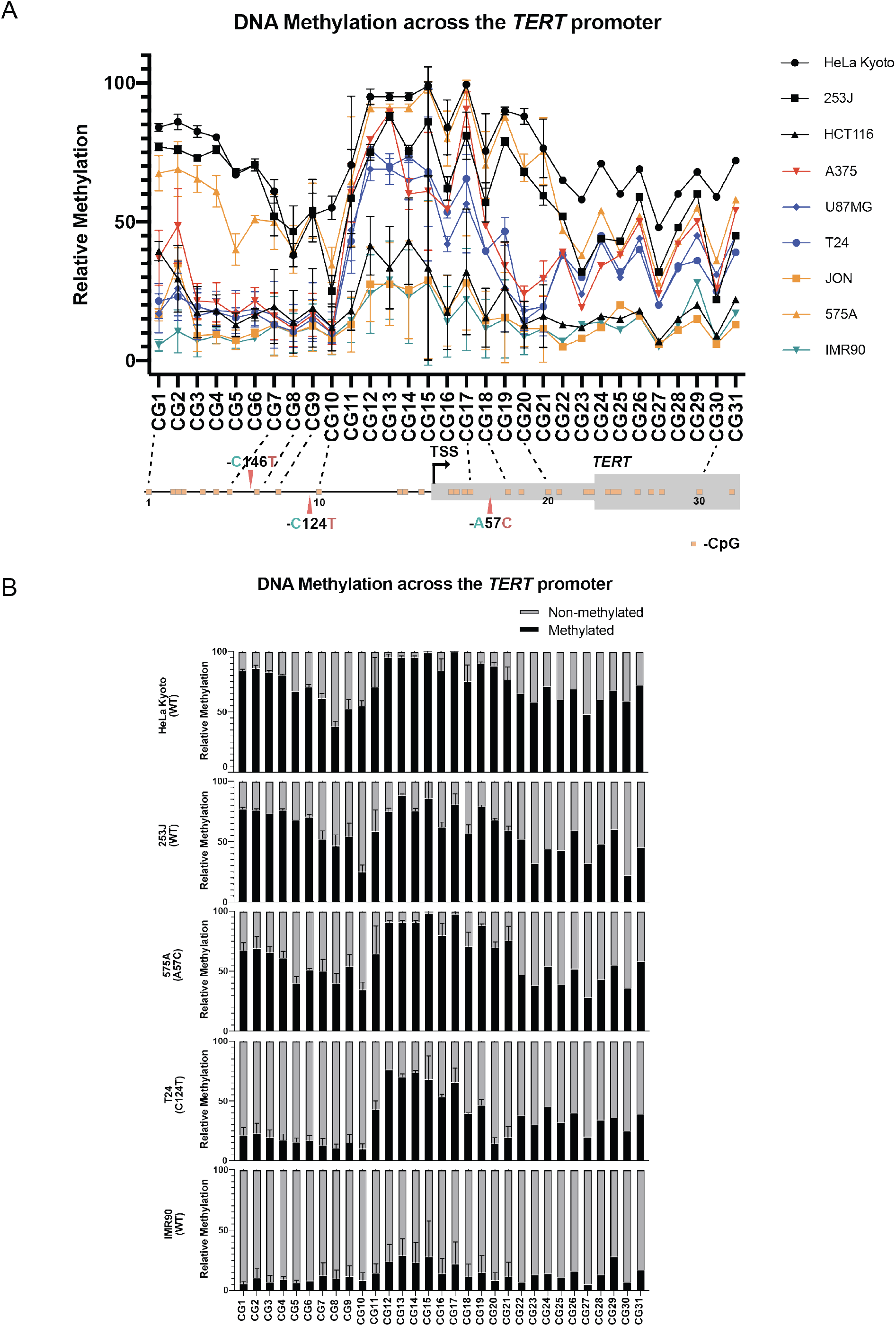
**(A)** Relative DNA methylation frequency of CpGs across the *TERT* promoter in cell lines with the WT promoter (HeLa Kyoto, 253J, HCT116, T24 & IMR90), the C146T mutation (A375), the C124T mutation (U87MG) or the A57C mutation (575A & JON). **(B)** DNA methylation on CpGs across the *TERT* promoter in selected cell lines. Generally, telomerase-positive cell lines exhibit higher methylation levels compared to primary fibroblasts IMR90.

**Supplementary Figure S3.**
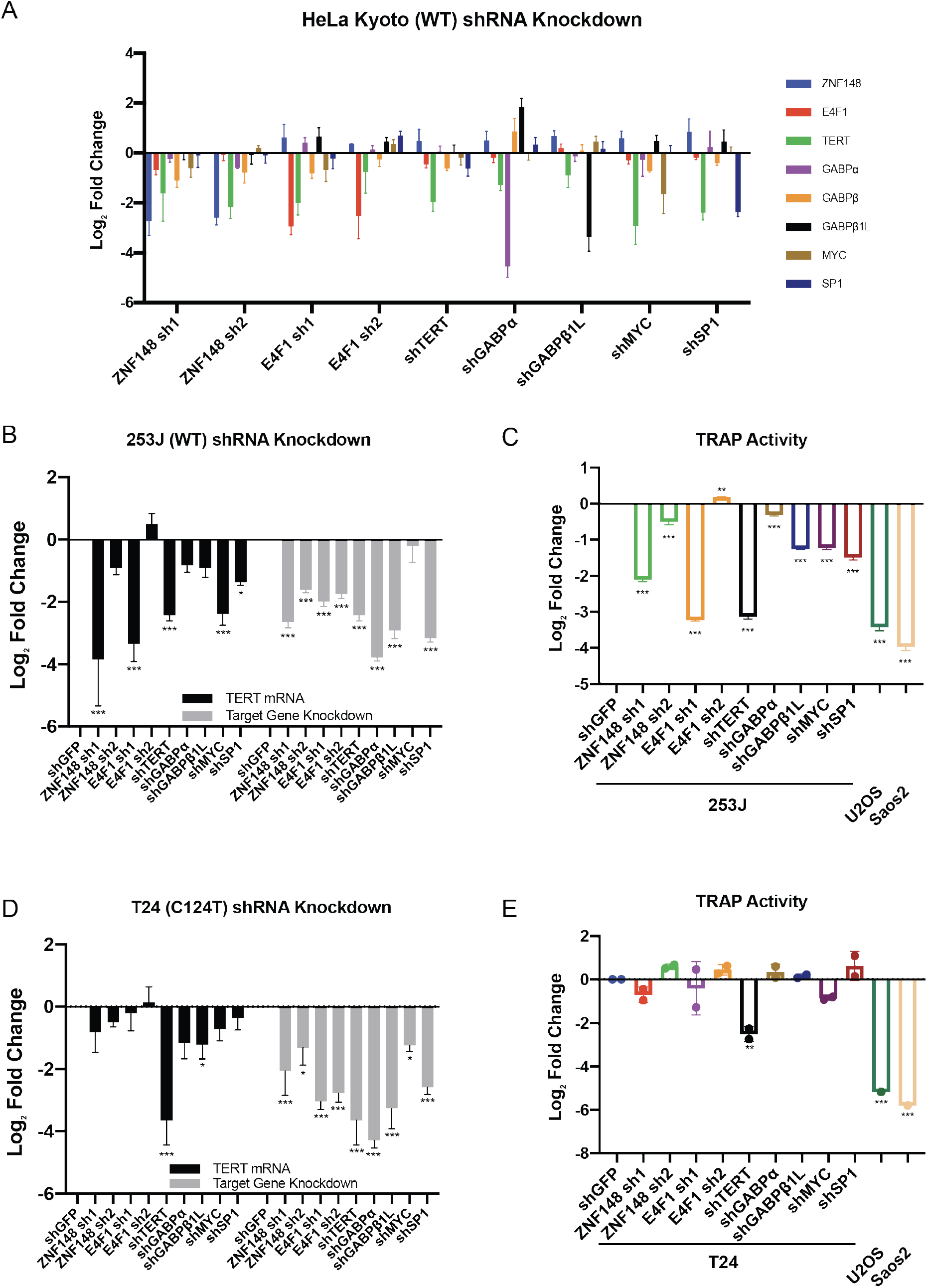
**(A)** mRNA expression data of ZNF148, E4F1, TERT, GABPα, GABPβ1L, MYC and Sp1 following 48+72 h post-shRNA knockdown in HeLa (48h virus transduction, 72 hours puromycin selection), with shGFP as control. Data shown as mean of values from three biological replicates. **(B)** mRNA expression data of *TERT* and target gene following 48+72 h post-shRNA knockdown in 253J, with shGFP as control. **(C)** TRAP assay measuring telomerase activity following 48+72 h post-shRNA knockdown in 253J (48h virus transduction, 72 hours puromycin selection), with shGFP as control. Telomerase-negative U2OS and Saos2 were used as negative controls. **(D)** mRNA expression data of *TERT* and target genes following 48+72 h post-shRNA knockdown in T24 (48h virus transduction, 72 hours puromycin selection), with shGFP as control. Data shown as mean of values from three biological replicates. **(E)** ddPCR-TRAP assay measuring telomerase activity following 48+72 h post-shRNA knockdown in T24, with shGFP as control. Telomerase-negative U2OS and Saos2 were used as negative controls. Data shown as mean of values from two biological replicates. All statistical significance was calculated using a two-sampled t-test, and the degree of significance is indicated as: * for p <0.05; ** for p <0.01; *** for p <0.001.

## Supplementary Tables

**Supplementary Table S1.**
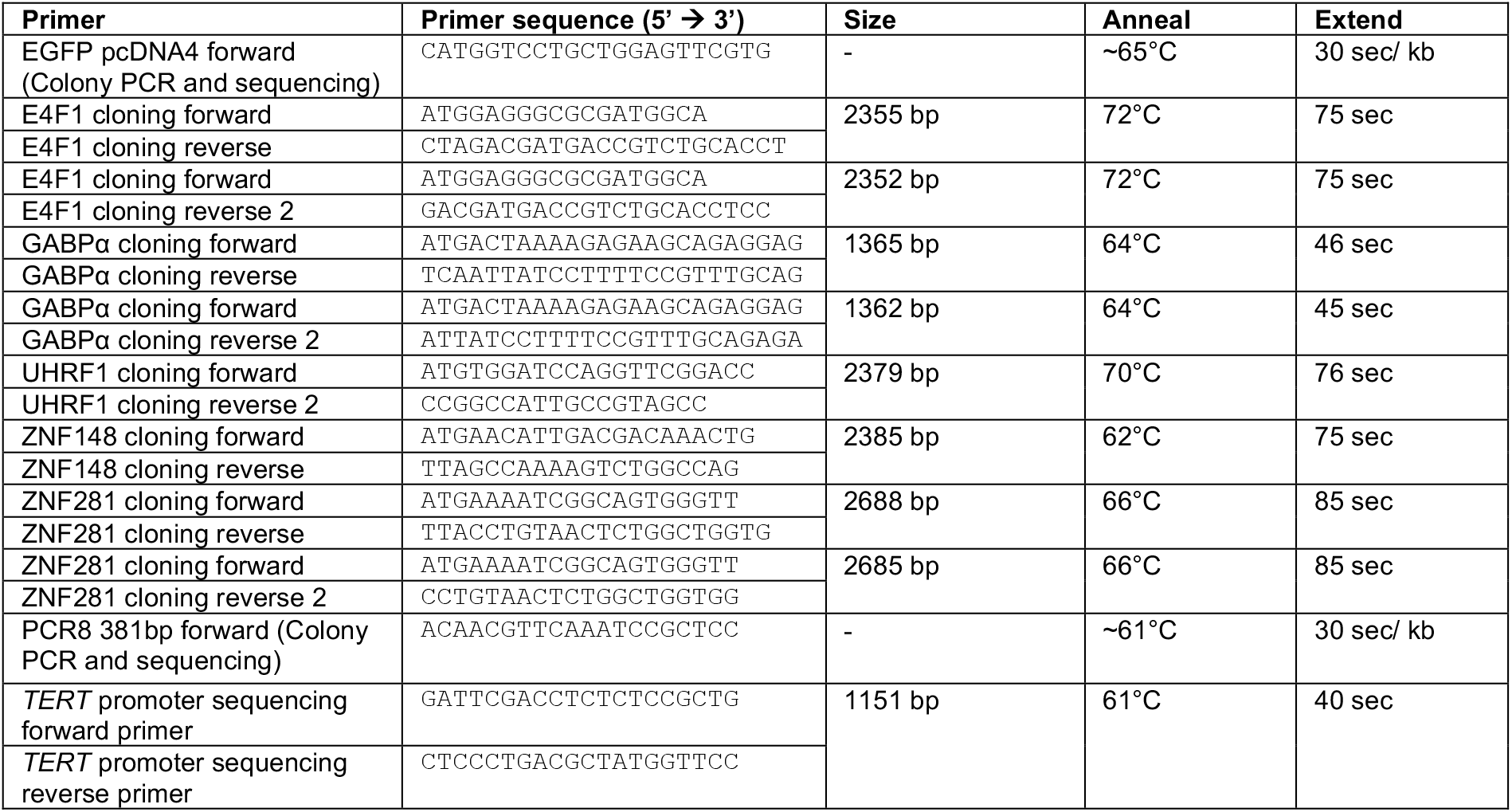
List of oligonucleotides used for cloning and sequencing of cDNA and gDNA sequences.

**Supplementary Table S2.**
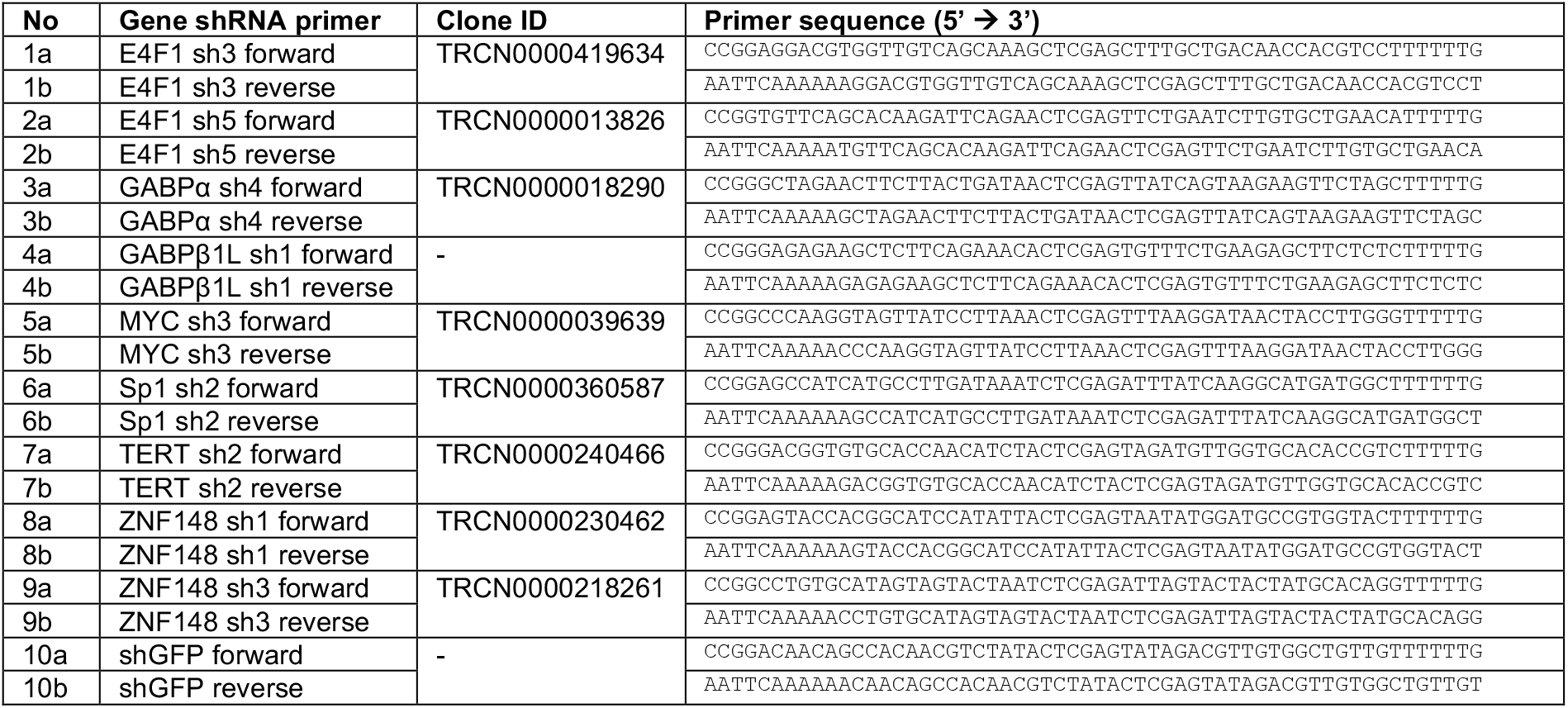
List of oligonucleotides used for shRNA generation.

**Supplementary Table S3.**
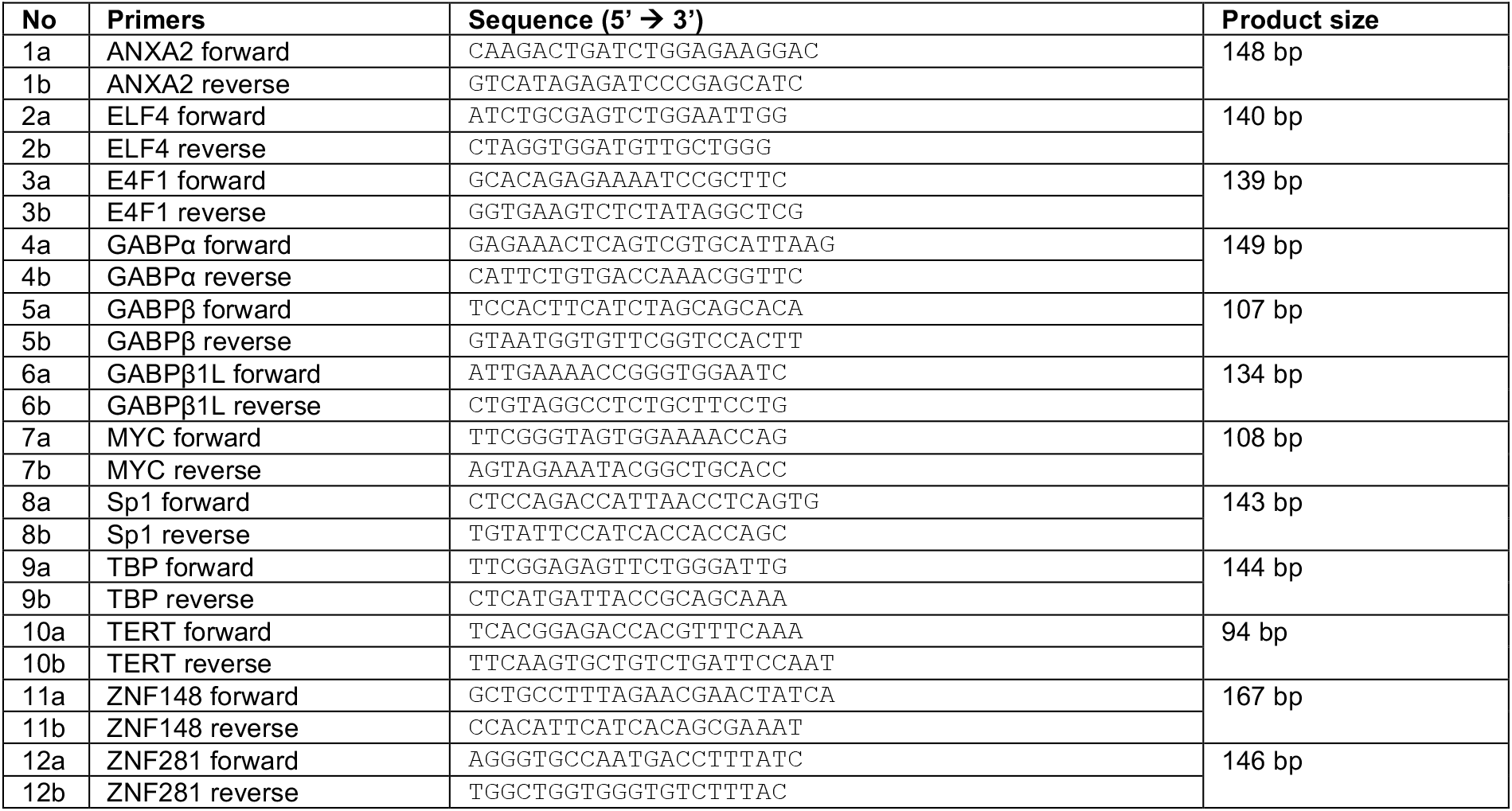
List of oligonucleotides used for quantiative PCR.

**Supplementary Table S4.**
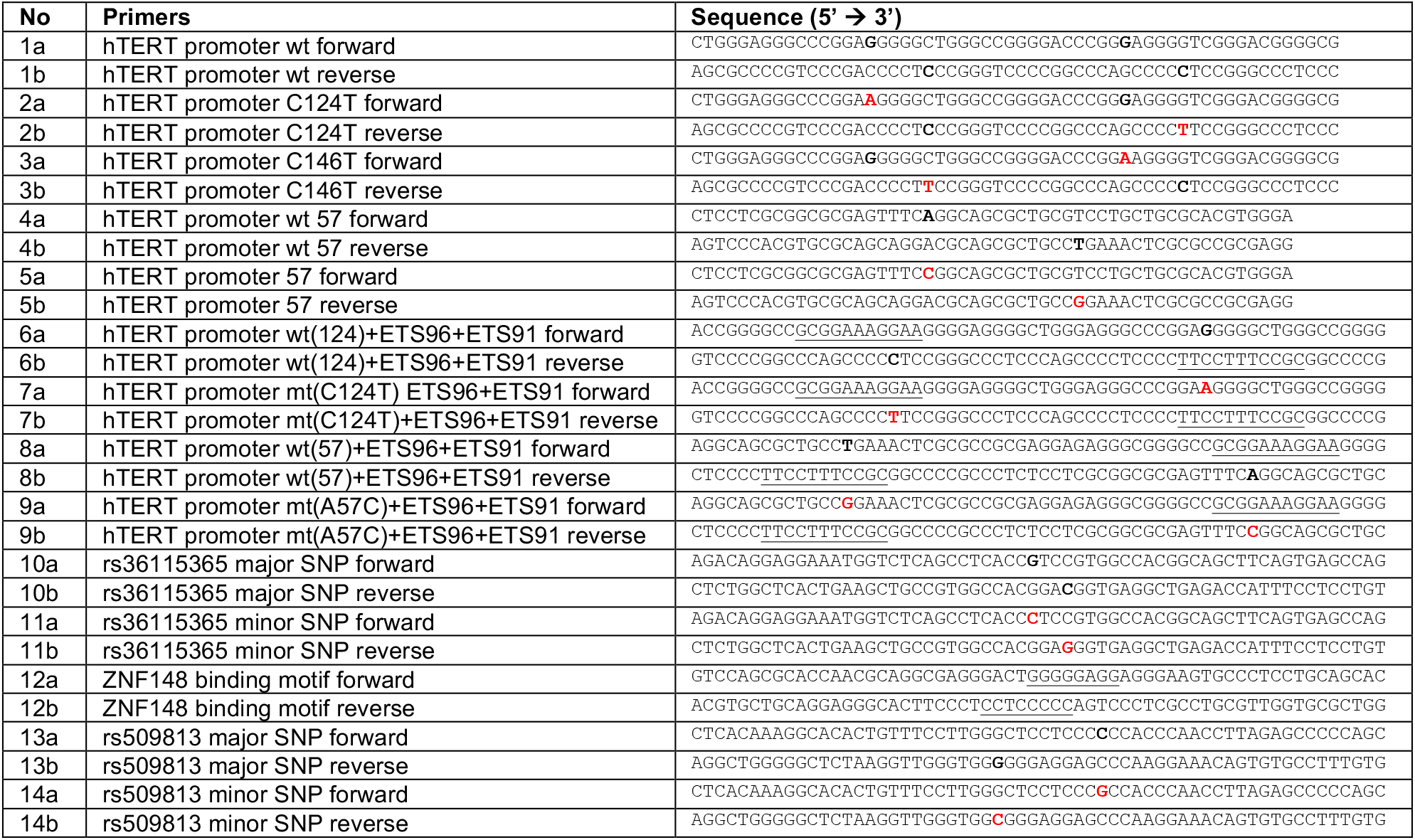
List of oligonucleotides used for DNA pulldown.

**Supplementary Table S5.** List of antibodies used for Western blot.

| Antibody | Company | Catalogue No. | Origin | Dilution used for WB |
| --- | --- | --- | --- | --- |
| α-GFP | Roche | 11814460001 | mouse monoclonal | 1:4000 |
| c-Myc | Santa-Cruz | sc-40 | mouse monoclonal | 1:250 |
| DM1 α-tubulin | MPI-CPG |  | mouse monoclonal | 1:10000 |
| E4F1 | Santa-Cruz | sc-514718 | mouse monoclonal | 1:500 |
| GABPα | Santa-Cruz | sc-28312 | mouse monoclonal | 1:250 |
| GABP-β1/2 | Santa-Cruz | sc-271571 | mouse monoclonal | 1:250 |
| GAPDH | Novus Biologicals | NB300-221 | Mouse monoclonal | 1:1000 |
| Sp1 | Santa-Cruz | sc-17824 | mouse monoclonal | 1:250 |
| ZNF148 | Santa-Cruz | sc-137171 | mouse monoclonal | 1:500 |

**Supplementary Table S6.**
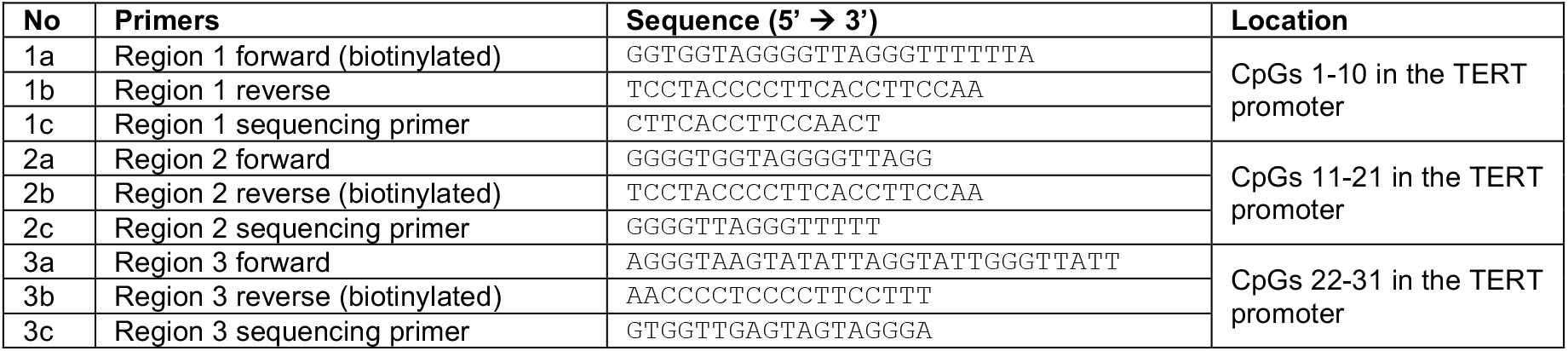
List of oligonucleotides used for pyrosequencing.

**Supplementary Table S7.** *TERT* promoter mutation status at different positions of various cell lines.

| No | Primers | -146 | -124 | -57 |
| --- | --- | --- | --- | --- |
| 1 | HeLa Kyoto | 100% C | 100% C | 100% A |
| 2 | HCT116 | 100% C | 100% C | 100% A |
| 3 | 253J | 100% C | 100% C | 100% A |
| 4 | A375 | 20% C, 80% T | 100% C | 100% A |
| 5 | U87MG | 100% C | 15% C, 85% T | 100% A |
| 6 | T24 | 100% C | 100% T | 100% A |
| 7 | JON | 100% C | 100% C | 40% A, 60% C |
| 8 | 575A | 100% C | 100% C | 80% A, 20% C |

**Supplementary Table S8.** MS data for SILAC-based in-vitro DNA reconstitution pull-downs using U87MG nuclear extract with C124T vs. WT TERT promoter probes.

**Supplementary Table S9.** MS data for SILAC-based in-vitro DNA reconstitution pull-downs using U87MG nuclear extract with C146T vs. WT TERT promoter probes.

**Supplementary Table S10.** MS data for SILAC-based in-vitro DNA reconstitution pull-downs using U87MG nuclear extract with A57C vs. WT TERT promoter probes.

## REFERENCES

1. Horn, S., Figl, A., Rachakonda, P.S., Fischer, C., Sucker, A., Gast, A., Kadel, S., Moll, I., Nagore, E., Hemminki, K., et al. (2013) TERT Promoter Mutations in Familial and Sporadic Melanoma. Science, 339, 959–961.

2. Huang, F.W., Hodis, E., Xu, M.J., Kryukov, G.V., Chin, L. and Garraway, L.A. (2013) Highly Recurrent TERT Promoter Mutations in Human Melanoma. Science, 339, 957–959.

3. Vinagre, J., Almeida, A., Pópulo, H., Batista, R., Lyra, J., Pinto, V., Coelho, R., Celestino, R., Prazeres, H., Lima, L., et al. (2013) Frequency of TERT promoter mutations in human cancers. Nat Commun, 4, 2185.

4. Borah, S., Xi, L., Zaug, A.J., Powell, N.M., Dancik, G.M., Cohen, S.B., Costello, J.C., Theodorescu, D. and Cech, T.R. (2015) Cancer. TERT promoter mutations and telomerase reactivation in urothelial cancer. Sci New York N Y, 347, 1006–10.

5. Killela, P.J., Reitman, Z.J., Jiao, Y., Bettegowda, C., Agrawal, N., Diaz, L.A., Friedman, A.H., Friedman, H., Gallia, G.L., Giovanella, B.C., et al. (2013) TERT promoter mutations occur frequently in gliomas and a subset of tumors derived from cells with low rates of self-renewal. Proc National Acad Sci, 110, 6021–6026.

6. Bell, R.J.A., Rube, H.T., Xavier-Magalhães, A., Costa, B.M., Mancini, A., Song, J.S. and Costello, J.F. (2016) Understanding TERT Promoter Mutations: A Common Path to Immortality. Mol Cancer Res, 14, 315–323.

7. Olivier, M., Hollstein, M. and Hainaut, P. (2010) TP53 Mutations in Human Cancers: Origins, Consequences, and Clinical Use. Csh Perspect Biol, 2, a001008.

8. Hurst, C.D., Platt, F.M. and Knowles, M.A. (2014) Comprehensive Mutation Analysis of the TERT Promoter in Bladder Cancer and Detection of Mutations in Voided Urine. Eur Urol, 65, 367–369.

9. Kinde, I., Munari, E., Faraj, S.F., Hruban, R.H., Schoenberg, M., Bivalacqua, T., Allaf, M., Springer, S., Wang, Y., Diaz, L.A., et al. (2013) TERT Promoter Mutations Occur Early in Urothelial Neoplasia and Are Biomarkers of Early Disease and Disease Recurrence in Urine. Cancer Res, 73, 7162– 7167.

10. Bell, R.J.A., Rube, H.T., Kreig, A., Mancini, A., Fouse, S.D., Nagarajan, R.P., Choi, S., Hong, C., He, D., Pekmezci, M., et al. (2015) Cancer. The transcription factor GABP selectively binds and activates the mutant TERT promoter in cancer. Sci New York N Y, 348, 1036–9.

11. Shay, J.W. and Bacchetti, S. (1997) A survey of telomerase activity in human cancer. Eur J Cancer, 33, 787–791.

12. Hanahan, D. and Weinberg, R.A. (2011) Hallmarks of Cancer: The Next Generation. Cell, 144, 646–674.

13. Moyzis, R.K., Buckingham, J.M., Cram, L.S., Dani, M., Deaven, L.L., Jones, M.D., Meyne, J., Ratliff, R.L. and Wu, J.R. (1988) A highly conserved repetitive DNA sequence, (TTAGGG)n, present at the telomeres of human chromosomes. Proc National Acad Sci, 85, 6622–6626.

14. Palm, W. and Lange, T. de (2008) How Shelterin Protects Mammalian Telomeres. Annu Rev Genet, 42, 301–334.

15. Olovnikov, A.M. (1971) [Principle of marginotomy in template synthesis of polynucleotides]. Dokl Akad Nauk Sssr+, 201, 1496–9.

16. Watson, J.D. (1972) Origin of Concatemeric T7DNA. Nat New Biology, 239, 197–201.

17. O’Sullivan, R.J. and Karlseder, J. (2010) Telomeres: protecting chromosomes against genome instability. Nat Rev Mol Cell Bio, 11, 171–181.

18. Wu, P., Takai, H. and de Lange, T. (2012) Telomeric 3′ Overhangs Derive from Resection by Exo1 and Apollo and Fill-In by POT1b-Associated CST. Cell, 150, 39–52.

19. Zhao, Y., Sfeir, A.J., Zou, Y., Buseman, C.M., Chow, T.T., Shay, J.W. and Wright, W.E. (2009) Telomere Extension Occurs at Most Chromosome Ends and Is Uncoupled from Fill-In in Human Cancer Cells. Cell, 138, 463–475.

20. Bonetti, D., Martina, M., Falcettoni, M. and Longhese, M.P. (2013) Telomere-end processing: mechanisms and regulation. Chromosoma, 123, 57–66.

21. Shay, J.W. and Wright, W.E. (2011) Role of telomeres and telomerase in cancer. Semin Cancer Biol, 21, 349–353.

22. Hunter, S.A., Iwei, Y., Ivanka, K., Aravindhan, S., Eric, T., Alexander, G., Reinhard, D., Jeffrey, N., Laura, P., Beth, R., et al. (2015) The Genetic Evolution of Melanoma from Precursor Lesions. New Engl J Med, 373, 1926–1936.

23. Lee, J.H., Lee, J.E., Kahng, J.Y., Kim, S.H., Park, J.S., Yoon, S.J., Um, J.-Y., Kim, W.K., Lee, J.-K., Park, J., et al. (2017) Human glioblastoma arises from subventricular zone cells with low-level driver mutations. Nature, 560, 243–247.

24. Nault, J.C., Mallet, M., Pilati, C., Calderaro, J., Bioulac-Sage, P., Laurent, C., Laurent, A., Cherqui, D., Balabaud, C., Zucman-Rossi, J., et al. (2013) High frequency of telomerase reverse-transcriptase promoter somatic mutations in hepatocellular carcinoma and preneoplastic lesions. Nat Commun, 4, 2218.

25. Lorbeer, F.K. and Hockemeyer, D. (2019) TERT promoter mutations and telomeres during tumorigenesis. Curr Opin Genet Dev, 60, 56–62.

26. Chiba, K., Lorbeer, F.K., Shain, A.H., McSwiggen, D.T., Schruf, E., Oh, A., Ryu, J., Darzacq, X., Bastian, B.C. and Hockemeyer, D. (2017) Mutations in the promoter of the telomerase gene TERT contribute to tumorigenesis by a two-step mechanism. Science, 357, 1416–1420.

27. Casillas, M.A., Brotherton, S.L., Andrews, L.G., Ruppert, J.M. and Tollefsbol, T.O. (2003) Induction of endogenous telomerase (hTERT) by c-Myc in WI-38 fibroblasts transformed with specific genetic elements. Gene, 316, 57–65.

28. Ge, Z., Li, W., Wang, N., Liu, C., Zhu, Q., Björkholm, M., Gruber, A. and Xu, D. (2010) Chromatin remodeling: recruitment of histone demethylase RBP2 by Madl for transcriptional repression of a Myc target gene, telomerase reverse transcriptase. Faseb J, 24, 579–586.

29. Wu, K.-J., Grandori, C., Amacker, M., Simon-Vermot, N., Polack, A., Lingner, J. and Dalla-Favera, R. (1999) Direct activation of TERT transcription by c-MYC. Nat Genet, 21, 220–224.

30. Kyo, S., Takakura, M., Fujiwara, T. and Inoue, M. (2008) Understanding and exploiting hTERT promoter regulation for diagnosis and treatment of human cancers. Cancer Sci, 99, 1528–38.

31. Liu, C., Fang, X., Ge, Z., Jalink, M., Kyo, S., Björkholm, M., Gruber, A., Sjöberg, J. and Xu, D. (2007) The Telomerase Reverse Transcriptase (hTERT) Gene Is a Direct Target of the Histone Methyltransferase SMYD3. Cancer Res, 67, 2626–2631.

32. Rajagopalan, D., Pandey, A.K., Xiuzhen, M.C., Lee, K.K., Hora, S., Zhang, Y., Chua, B.H., Kwok, H.S., Bhatia, S.S., Deng, L.W., et al. (2017) TIP60 represses telomerase expression by inhibiting Sp1 binding to the TERT promoter. Plos Pathog, 13, e1006681.

33. Dessain, S.K., Yu, H., Reddel, R.R., Beijersbergen, R.L. and Weinberg, R.A. (2000) Methylation of the human telomerase gene CpG island. Cancer Res, 60, 537–41.

34. Renaud, S., Loukinov, D., Abdullaev, Z., Guilleret, I., Bosman, F.T., Lobanenkov, V. and Benhattar, J. (2007) Dual role of DNA methylation inside and outside of CTCF-binding regions in the transcriptional regulation of the telomerase hTERT gene. Nucleic Acids Res, 35, 1245–1256.

35. Stern, J.L., Paucek, R.D., Huang, F.W., Ghandi, M., Nwumeh, R., Costello, J.C. and Cech, T.R. (2017) Allele-Specific DNA Methylation and Its Interplay with Repressive Histone Marks at Promoter-Mutant TERT Genes. Cell Reports, 21, 3700–3707.

36. Choi, J.-H., Min, N.Y., Park, J., Kim, J.H., Park, S.H., Ko, Y.J., Kang, Y., Moon, Y.J., Rhee, S., Ham, S.W., et al. (2010) TSA-induced DNMT1 down-regulation represses hTERT expression via recruiting CTCF into demethylated core promoter region of hTERT in HCT116. Biochem Bioph Res Co, 391, 449–454.

37. Esopi, D., Graham, M.K., Brosnan-Cashman, J.A., Meyers, J., Vaghasia, A., Gupta, A., Kumar, B., Haffner, M.C., Heaphy, C.M., Marzo, A.M.D., et al. (2020) Pervasive promoter hypermethylation of silenced TERT alleles in human cancers. Cell Oncol, 43, 847–861.

38. Stern, J.L., Theodorescu, D., Vogelstein, B., Papadopoulos, N. and Cech, T.R. (2015) Mutation of the TERT promoter, switch to active chromatin, and monoallelic TERT expression in multiple cancers. Gene Dev, 29, 2219–2224.

39. Wang, Z. and Zhang, Q. (2009) Genome-Wide Identification and Evolutionary Analysis of the Animal Specific ETS Transcription Factor Family. Evol Bioinform, 5, EBO.S2948.

40. Hollenhorst, P.C., Shah, A.A., Hopkins, C. and Graves, B.J. (2007) Genome-wide analyses reveal properties of redundant and specific promoter occupancy within the ETS gene family. Gene Dev, 21, 1882–1894.

41. Shore, P. and Sharrocks, A.D. (1995) The ETS-domain transcription factors Elk-1 and SAP-1 exhibit differential DNA binding specifitoies. Nucleic Acids Res, 23, 4698–4706.

42. Oikawa, T. and Yamada, T. (2003) Molecular biology of the Ets family of transcription factors. Gene, 303, 11–34.

43. Mancini, A., Xavier-Magalhães, A., Woods, W.S., Nguyen, K.-T., Amen, A.M., Hayes, J.L., Fellmann, C., Gapinske, M., McKinney, A.M., Hong, C., et al. (2018) Disruption of the β1L Isoform of GABP Reverses Glioblastoma Replicative Immortality in a TERT Promoter Mutation-Dependent Manner. Cancer Cell, 34, 513–528.e8.

44. Li, Y., Zhou, Q.-L., Sun, W., Chandrasekharan, P., Cheng, H.S., Ying, Z., Lakshmanan, M., Raju, A., Tenen, D.G., Cheng, S.-Y., et al. (2015) Non-canonical NF-κB signalling and ETS1/2 cooperatively drive C250T mutant TERT promoter activation. Nat Cell Biol, 17, 1327–1338.

45. Vallarelli, A.F., Rachakonda, P.S., André, J., Heidenreich, B., Riffaud, L., Bensussan, A., Kumar, R. and Dumaz, N. (2016) TERT promoter mutations in melanoma render TERT expression dependent on MAPK pathway activation. Oncotarget, 7, 53127–53136.

46. Cristofari, G. and Lingner, J. (2006) Telomere length homeostasis requires that telomerase levels are limiting. Embo J, 25, 565–574.

47. Kappei, D., Scheibe, M., Paszkowski-Rogacz, M., Bluhm, A., Gossmann, T.I., Dietz, S., Dejung, M., Herlyn, H., Buchholz, F., Mann, M., et al. (2017) Phylointeractomics reconstructs functional evolution of protein binding. Nat Commun, 8, 14334.

48. Cox, J. and Mann, M. (2008) MaxQuant enables high peptide identification rates, individualized p.p.b.-range mass accuracies and proteome-wide protein quantification. Nat Biotechnol, 26, 1367– 1372.

49. Makowski, M.M., Willems, E., Fang, J., Choi, J., Zhang, T., Jansen, P.W.T.C., Brown, K.M. and Vermeulen, M. (2016) An interaction proteomics survey of transcription factor binding at recurrent TERT promoter mutations. Proteomics, 16, 417–426.

50. Mondal, S., Ramanathan, M., Miao, W., Meyers, R.M., Rao, D., Lopez-Pajares, V., Siprashvili, Z., Reynolds, D.L., Porter, D.F., Ferguson, I., et al. (2022) PROBER identifies proteins associated with programmable sequence-specific DNA in living cells. Nat Methods, 19, 959–968.

51. Fang, J., Jia, J., Makowski, M., Xu, M., Wang, Z., Zhang, T., Hoskins, J.W., Choi, J., Han, Y., Zhang, M., et al. (2017) Functional characterization of a multi-cancer risk locus on chr5p15.33 reveals regulation of TERT by ZNF148. Nat Commun, 8, 15034.

52. Butter, F., Davison, L., Viturawong, T., Scheibe, M., Vermeulen, M., Todd, J.A. and Mann, M. (2012) Proteome-Wide Analysis of Disease-Associated SNPs That Show Allele-Specific Transcription Factor Binding. Plos Genet, 8, e1002982.

53. Cam, L.L., Linares, L.K., Paul, C., Julien, E., Lacroix, M., Hatchi, E., Triboulet, R., Bossis, G., Shmueli, A., Rodriguez, M.S., et al. (2006) E4F1 Is an Atypical Ubiquitin Ligase that Modulates p53 Effector Functions Independently of Degradation. Cell, 127, 775–788.

54. Rooney, R.J., Rothammer, K. and Fernandes, E.R. (1998) Mutational analysis of p50E4F suggests that DNA binding activity is mediated through an alternative structure in a zinc finger domain that is regulated by phosphorylation. Nucleic Acids Res, 26, 1681–1688.

55. Xuan Lin, Q.X., Sian, S., An, O., Thieffry, D., Jha, S. and Benoukraf, T. (2018) MethMotif: an integrative cell specific database of transcription factor binding motifs coupled with DNA methylation profiles. Nucleic Acids Res, 47, gky1005.

56. Cong, Y.-S., Wen, J. and Bacchetti, S. (1999) The Human Telomerase Catalytic Subunit hTERT: Organization of the Gene and Characterization of the Promoter. Hum Mol Genet, 8, 137–142.

57. Horikawa, I., Cable, P.L., Afshari, C. and Barrett, J.C. (1999) Cloning and characterization of the promoter region of human telomerase reverse transcriptase gene. Cancer Res, 59, 826–30.

58. Takakura, M., Kyo, S., Kanaya, T., Hirano, H., Takeda, J., Yutsudo, M. and Inoue, M. (1999) Cloning of human telomerase catalytic subunit (hTERT) gene promoter and identification of proximal core promoter sequences essential for transcriptional activation in immortalized and cancer cells. Cancer Res, 59, 551–7.

59. Cong, Y.-S. and Bacchetti, S. (2000) Histone Deacetylation Is Involved in the Transcriptional Repression of hTERT in Normal Human Cells*. J Biol Chem, 275, 35665–35668.

60. Hou, M., Wang, X., Popov, N., Zhang, A., Zhao, X., Zhou, R., Zetterberg, A., Björkholm, M., Henriksson, M., Gruber, A., et al. (2002) The Histone Deacetylase Inhibitor Trichostatin A Derepresses the Telomerase Reverse Transcriptase (hTERT) Gene in Human Cells. Exp Cell Res, 274, 25–34.

61. Takakura, M., Kyo, S., Sowa, Y., Wang, Z., Yatabe, N., Maida, Y., Tanaka, M. and Inoue, M. (2001) Telomerase activation by histone deacetylase inhibitor in normal cells. Nucleic Acids Res, 29, 3006–3011.

62. Ludlow, A.T., Robin, J.D., Sayed, M., Litterst, C.M., Shelton, D.N., Shay, J.W. and Wright, W.E. (2014) Quantitative telomerase enzyme activity determination using droplet digital PCR with single cell resolution. Nucleic Acids Res, 42, e104–e104.

63. Ludlow, A.T., Shelton, D., Wright, W.E. and Shay, J.W. (2018) Digital PCR, Methods and Protocols. Methods Mol Biology Clifton N J, 1768, 513–529.

64. Wei, G., Badis, G., Berger, M.F., Kivioja, T., Palin, K., Enge, M., Bonke, M., Jolma, A., Varjosalo, M., Gehrke, A.R., et al. (2010) Genome-wide analysis of ETS-family DNA-binding in vitro and in vivo. Embo J, 29, 2147–2160.

65. Fernandes, E.R. and Rooney, R.J. (1997) The adenovirus E1A-regulated transcription factor E4F is generated from the human homolog of nuclear factor phiAP3. Mol Cell Biol, 17, 1890–1903.

66. Lee, K.A. and Green, M.R. (1987) A cellular transcription factor E4F1 interacts with an E1a-inducible enhancer and mediates constitutive enhancer function in vitro. Embo J, 6, 1345–1353.

67. Rodier, G., Kirsh, O., Baraibar, M., Houlès, T., Lacroix, M., Delpech, H., Hatchi, E., Arnould, S., Severac, D., Dubois, E., et al. (2015) The Transcription Factor E4F1 Coordinates CHK1-Dependent Checkpoint and Mitochondrial Functions. Cell Reports, 11, 220–233.

68. Butter, F., Kappei, D., Buchholz, F., Vermeulen, M. and Mann, M. (2010) A domesticated transposon mediates the effects of a single-nucleotide polymorphism responsible for enhanced muscle growth. Embo Rep, 11, 305–311.

69. Liu, N.Q., Huurne, M. ter, Nguyen, L.N., Peng, T., Wang, S.-Y., Studd, J.B., Joshi, O., Ongen, H., Bramsen, J.B., Yan, J., et al. (2017) The non-coding variant rs1800734 enhances DCLK3 expression through long-range interaction and promotes colorectal cancer progression. Nat Commun, 8, 14418.

70. Perez-Riverol, Y., Bai, J., Bandla, C., García-Seisdedos, D., Hewapathirana, S., Kamatchinathan, S., Kundu, D.J., Prakash, A., Frericks-Zipper, A., Eisenacher, M., et al. (2021) The PRIDE database resources in 2022: a hub for mass spectrometry-based proteomics evidences. Nucleic Acids Res, 50, D543–D552.

